# Mechanism of negative membrane curvature generation by I-BAR domains

**DOI:** 10.1101/2020.08.19.256925

**Authors:** Binod Nepal, Aliasghar Sepehri, Themis Lazaridis

## Abstract

The membrane sculpting ability of BAR domains has been attributed to the intrinsic curvature of their banana-shaped dimeric structure. However, there is often a mismatch between this intrinsic curvature and the diameter of the membrane tubules generated. I-BAR domains have been especially mysterious: they are almost flat but generate high negative membrane curvature. Here, we use atomistic implicit-solvent computer modeling to show that the membrane bending of the IRSP53 I-BAR domain is dictated by its higher oligomeric structure, whose curvature is completely unrelated to the intrinsic curvature of the dimer. Two other I-BARs gave similar results, whereas a flat F-BAR sheet developed a concave membrane binding interface, consistent with its observed positive membrane curvature generation. Laterally interacting helical spirals of I-BAR dimers on tube interiors are stable and have an enhanced binding energy that is sufficient for membrane bending to experimentally observed tubule diameters at a reasonable surface density.

---

Remodeling of the cell membrane is involved in key biological processes such as endocytosis, viral budding, and membrane fission/fusion ^1-8^. This is usually accomplished by specialized proteins that can sense and generate membrane curvature. One important class of such proteins is the BAR domain family, consisting of crescent-shaped dimers and classified into three groups. Classical BAR/N-BAR domains generate or stabilize high, positive membrane curvature with intrinsic radius from 5.5 to 40 nm ^9-11,12^. The N-BAR domains contain an additional N-terminal amphipathic helix (H0 helix), which can assist in membrane binding and/or curvature generation. F-BAR domains tend to generate or stabilize lower positive membrane curvature than classical BAR domains with tubule diameters ranging from 64-113 nm ^13^. Finally, inverse BAR (I-BAR) domains stabilize or generate negative membrane curvature. These have a positively charged surface that is slightly convex in the crystal structures, which is thought to help them bind efficiently on the concave surface of the membrane. I-BAR domains are present in several proteins, including insulin receptor substrate P53 (IRSp53), insulin receptor tyrosine kinase receptor (IRTKS), missing-in-metastasis (MIM) and actin-bundling protein with BAIAP2 homology (ABBA). IRSp53 and IRTKS I-BAR domains bind to the membrane mainly through electrostatic interactions but MIM and ABBA also insert their N-terminal helix into the membrane interior ^14^.

There is evidence that generation of global membrane curvature requires assemblies of BAR proteins. For example, cryo-EM studies found that F-BAR domains polymerize into helical coats, stabilized by both tip-to-tip and lateral interactions ^13^. It is possible that I-BAR proteins behave similarly but evidence for this is currently lacking. The binding of these proteins to PI(4,5)P2 further complicates the mechanism, as I-BAR domains were found to cause PI(4,5)P2 clustering ^15,16^. Asymmetric distribution of PI(4,5)P2 promotes membrane curvature generation even without peptides ^17^ or just with simple cations like Ca^2+ 18^. Proteins that generate negative membrane curvature bind to the inner leaflet of tubules, and this makes them harder to study. Specialized techniques have been developed for negative curvature sensing peptides in the last five years. For example, pulling tubes from a GUV with encapsulated peptides provided a way to determine the preference of the IRSp53 I-BAR domain for ∼18 nm radius ^19^. More recently, a novel method utilizing protein sorting on tubular filopodia of varying diameter showed preferential binding to 25-nm and 19-nm radii for MIM I-BAR and IRSp53, respectively ^20^.

Computer simulations could provide insights but are limited by the system size and computational resources. Few studies have appeared on I-BAR domains. All-atom simulations usually consider one dimer on the membrane surface for a short amount of time ^21,22^, while coarse-grained simulations can handle large systems, at the cost of lower accuracy ^23,24^. Combining all-atom MD simulation and site mutation analysis showed that about 30 salt bridges are formed between I-BAR and the membrane head groups ^22^. Membrane deformation was explained in terms of PI(4,5)P2 lipid clustering caused by these salt bridges. A coarse-grained molecular dynamics (MD) simulation showed the generation of negative curvature but shallower than that generated experimentally ^21^. Moreover, the peptide was not binding through its convex surface and, depending on the binding orientation, it also generated positive membrane curvature. Another study showed similar I-BAR mediated PI(4,5)P2 clustering on the lower leaflet and curvature generation was postulated based on lipid asymmetry ^25^. The most recent coarse-grained MD simulation revealed that the peptide tends to orient along the axis of the 20 nm radius tube ^23^. This result is puzzling, considering the curvature preference for 20 nm tubes. If individual I-BAR domains do not prefer to be aligned with the curvature, then how can they sense or stabilize it?

One possible explanation is that I-BAR domains form higher oligomers that are responsible for curvature sensing and generation. If so, how does each I-BAR dimer arrange to form higher oligomers? IRSP53 I-BAR generates tubules with average diameter of about 43-nm ^14^. What dictates this? In the present work we sought answers to these questions using an implicit membrane model for curved membranes ^26^ that allows treatment of the proteins in atomistic detail.

## Results

### Binding orientation, energetics and curvature sensitivity of the dimer

Four simulations, each initiated from a different orientation with respect to the membrane, showed that the dimer always binds with a well-defined interface (Fig. 1a). This interface contains a large number of positively charged residues making varying contributions to binding (Table S1). Consistent with previous simulations and mutation studies ^21,22^, the residues near the end regions of the dimer interact more strongly with the membrane than those in the middle region. Surprisingly, due to the middle region, the binding interface appears not convex but concave. The dimer binds similarly onto the interior of a 30-nm spherical membrane (Fig. 1c). The same positively charged residues contribute to membrane binding but the fit appears improved. In the binding configuration of the dimer on the exterior 30-nm spherical surface (Fig. 1b) the middle region positive residues bind somewhat better, but there is significant loss of interactions in the arms. It seems that what matters for curvature sensing is not the curvature of the entire surface but only that of the two arms alone. The angle between the two arms (Fig. 1c) is flatter on the flat membrane compared to the interior of the spherical surface (Fig. S1).

**Figure 1.**
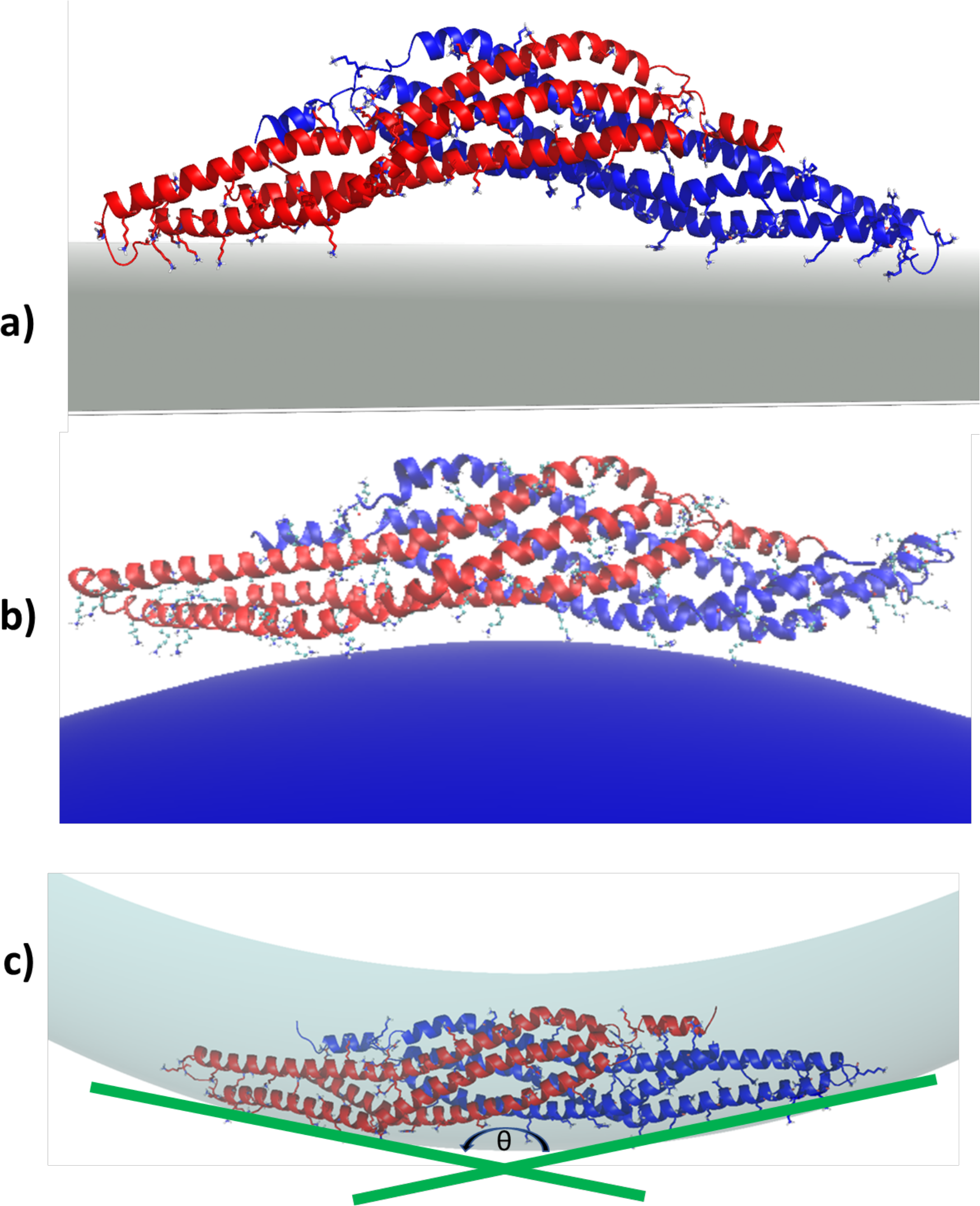
The binding configuration of the dimer on the flat membrane surface (a) and on a spherical surface of radius 30 nm from the outside (b) and from the inside (c), after 20 ns simulation. Transparent grey in (c) represents the surface of the inner leaflet of the vesicle on which the protein is sitting. The positively charged residues are shown in stick model. Shown is one of four independent trials, which give very similar results. θ is the angle between the axes of the helices defined by residues 121-147.

The orientational preference of the I-BAR dimer on cylindrical membrane tubes is presented in Table S2. In narrow tubes, inside or outside, the dimer tends to orient parallel to the tube axis, in agreement with recent coarse-grained simulations^23^. On the larger tubes this orientational preference persists only for the tube exterior, indicating that I-BAR dislikes especially a convex surface. The binding energies of the IRSP53 I-BAR dimer on 30% anionic spherical and cylindrical membrane surfaces are shown in Table 1. Binding energies on the interior spherical surface and on both cylindrical surfaces do not change much with curvature (the difference is within 0.3 kcal/mol). The latter is due to the preferred parallel orientation of the dimer to the tube axis. Only the exterior spherical surface is clearly less favorable at high curvature. Comparing the interior cylindrical and spherical surfaces, the cylindrical surface is slightly preferred, especially at higher curvature.

**Table 1.**
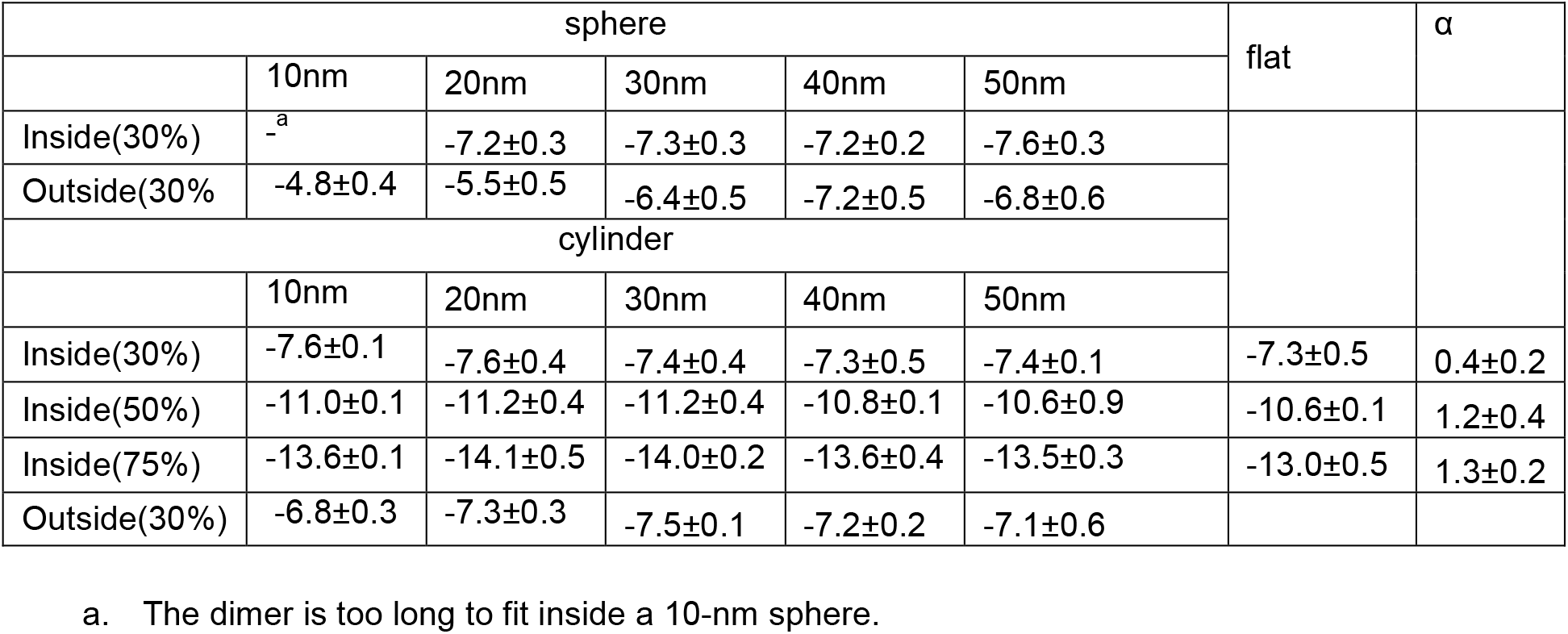
Binding energies (kcal/mol) of I-BAR dimer on cylindrical and spherical lipid membranes of different radius and anionic fraction as indicated. The curvature sensitivity parameter α is determined from the data for radii 20 nm to 50 nm.

The curvature sensitivities of the dimer were also determined at different anionic fractions (Table 1). At 30% anionic fraction, there is only a slight distinction between high curvature and the flat membrane surface. At higher anionic fractions, which could mimic the effect of anionic lipid clustering, binding energy is maximal at 20 nm radius and minimal on the flat surface. The curvature sensitivity parameter α increases significantly on increasing anionic fraction. The ratio of the dissociation constants for 20 nm radius to flat membrane increases from 1 to 3 to 6 on moving from anionic fraction 30% to 50% to 75%. Since I-BAR dimers align with the tube axis, the observed curvature sensitivity is attributed to the stronger electrostatic interaction in the narrow-sized tubes. The sorting ratio between the planar membrane and the 20-nm radius tubular membrane (Table S3) is 1.8 at 30% anionic membrane and increases to 6.4 upon increasing anionic percentage to 75%. A statistical analysis of these energy differences is shown in Table S4.

### Free simulations of multiple I-BAR dimers on the flat membrane surface

To identify possible oligomeric structures that might be responsible for generating or stabilizing tubular membranes, 2-20 I-BAR dimers were arranged on a rectangular lattice at a distance of 5 to 40 nm and were freely simulated on a flat membrane surface. Four independent runs with two dimers resulted in mostly lateral interactions having various alignments and stabilized by salt bridges. This is expected due to the presence of an almost uniform distribution of the positively and negatively charged residues throughout the length of the dimer (Fig. S2). Lateral interactions (Fig. 2b and 2c) were more probable than end-to-end interactions (Fig. 2d). Furthermore, the end-to-end interaction of just two dimers was not stable and eventually resulted in a laterally interacting dimer. When four dimers were freely simulated on the flat membrane surface, three of them came together and formed a laterally overlapping higher oligomer (Fig. 2e). In further simulations of 6, 8, 9 and 20 dimers on a flat membrane both lateral and end-to-end interactions were encountered, but lateral interactions were more pronounced (e.g. Fig. S3 for 20 dimers). The oligomerization interface obtained from the free simulations on the flat membrane (Fig. 2b) is similar to that observed in the crystal structure (PDB id 1Y2O).

**Figure 2.**
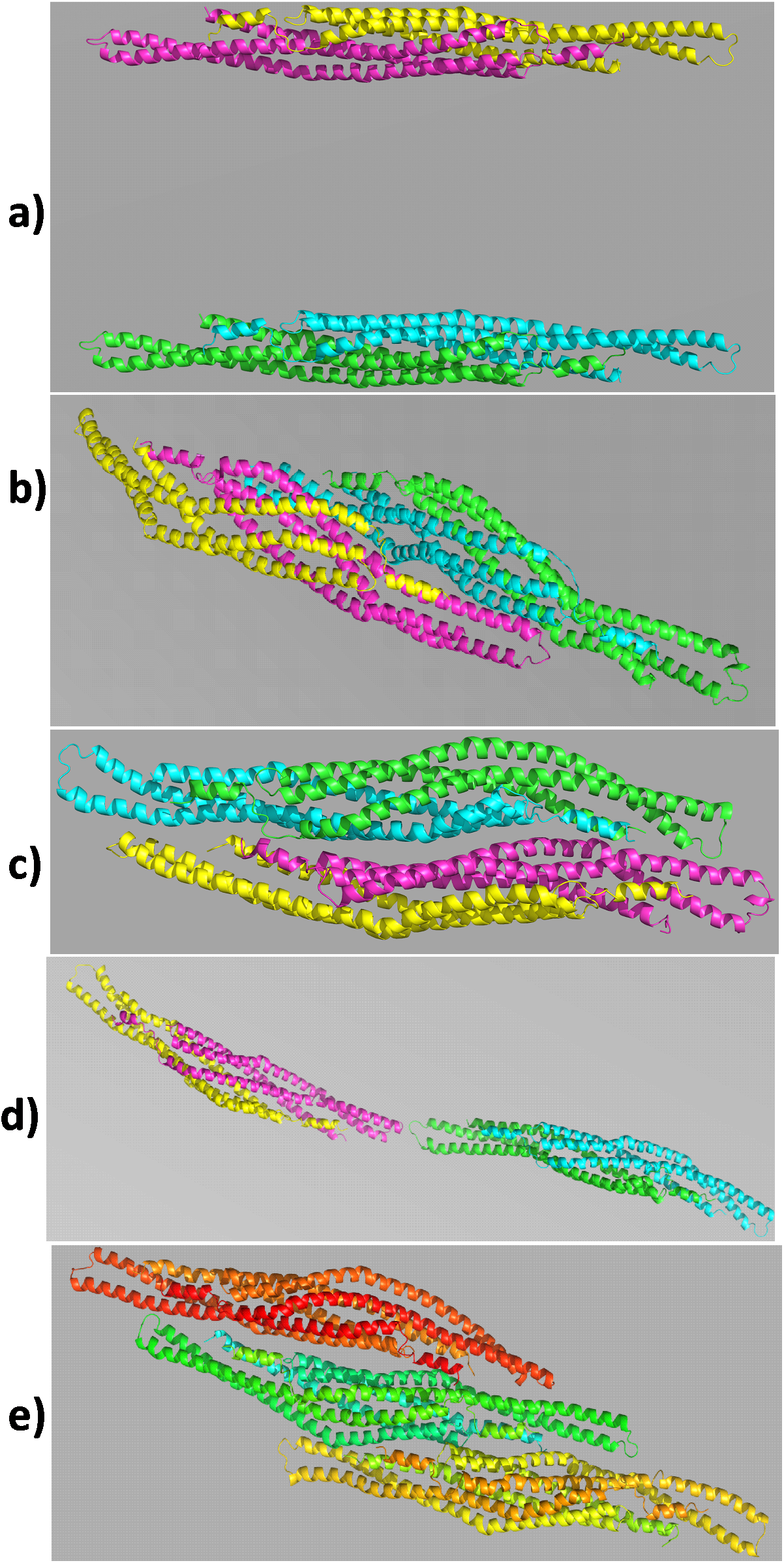
Initial (a) and final (b,c,d) configurations of the free simulations of two I-BAR dimers on the flat membrane surface. e) trimeric oligomer obtained from the free simulation of four dimers on the flat membrane surface. Top view is shown in all figures.

Free simulations of multiple I-BAR dimers were also carried out on the interior surface of a tubular membrane of 20-nm radius starting parallel or perpendicular to the tube axis. Again, the dimers interacted with each other both laterally and end-to-end. Most of the dimers tended to orient parallel to the tube axis. Hence, increasing the protein concentration to ∼544 dimers/μm^2^ does not lead to a preference for perpendicular orientation relative to the tube axis.

### Simulation of a chain and a sheet of dimers

Using the packing observed in the crystal structure and in the free simulations we generated a linear chain of dimers (Fig. 3a) and simulated it in implicit water and on a flat membrane surface. In water the chain bent into a spiral shape with radius close to 20 nm (Fig. 3b). Importantly, the membrane binding interface lies on the outer surface of the spiral, indicating that the spiral would readily bind to a concave surface. As mentioned earlier, in free simulations the lateral alignment between the dimers can vary. Regardless of the alignment, all constructed planar chains gave a spiral structure. On the flat implicit membrane surface the simulation resulted in a 2-dimensional bent structure (Fig. 3c).

**Figure 3.**
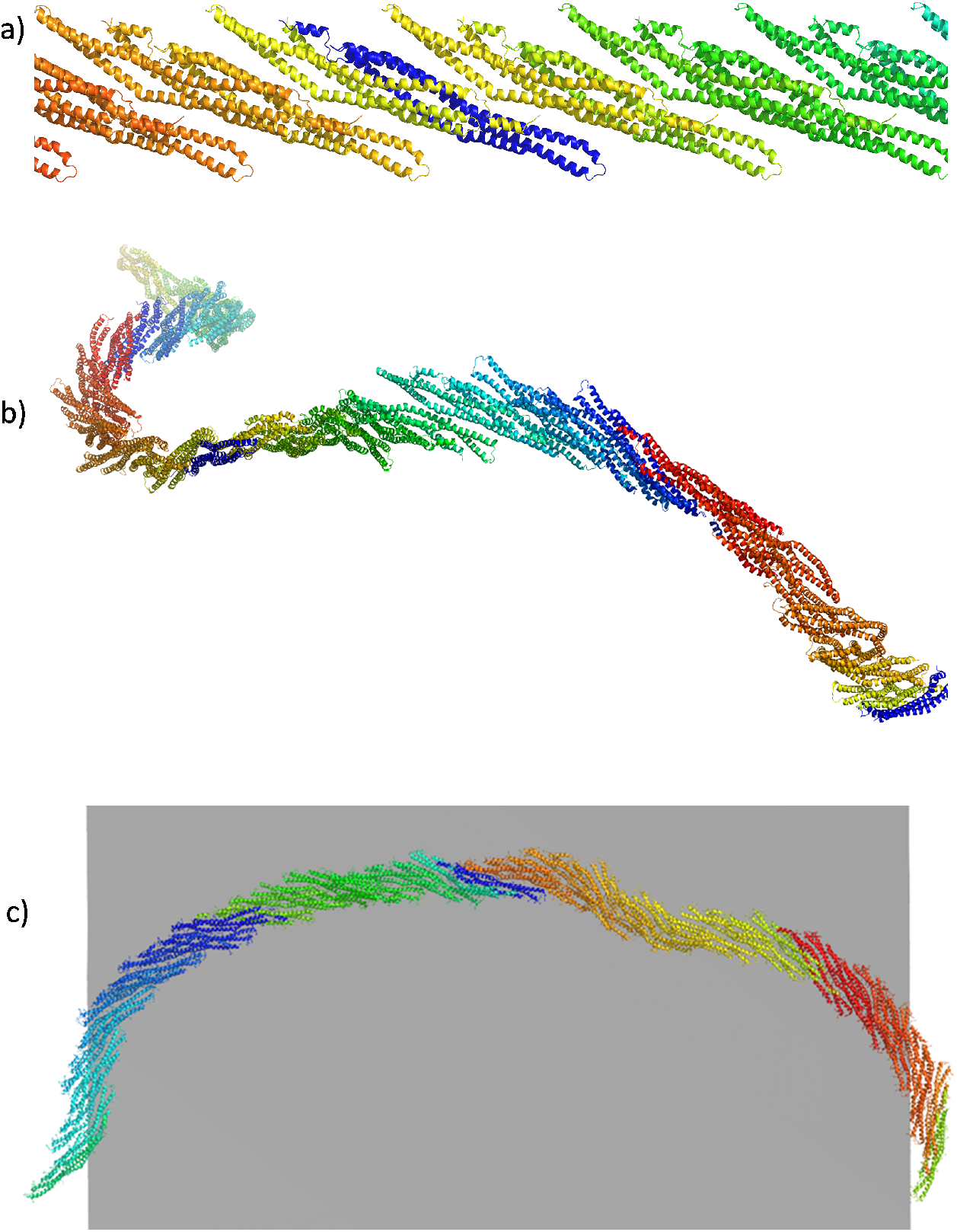
a) Initial arrangement of a chain of dimers based on the crystal structure, (b) final configuration obtained after simulation in implicit water, and c) final configuration obtained from the simulation on a flat membrane surface (view from above).

Similarly, a planar I-BAR sheet was constructed containing both lateral and end-to-end interactions (Fig. 4a) and was simulated in implicit water. The sheet rapidly turned into a tubular structure (Fig. 4b,c) whose membrane binding interface is on the outside, suitable for fitting into the interior of an anionic tubular membrane. Importantly, the sheet bent along the direction of the lateral overlap of the dimers, indicating that the bent structure of the I-BAR oligomer arises from the lateral interactions. The obtained structure had a radius of ∼13 nm, slightly lower than the preferred curvature determined by experiment ^14^.

**Figure 4.**
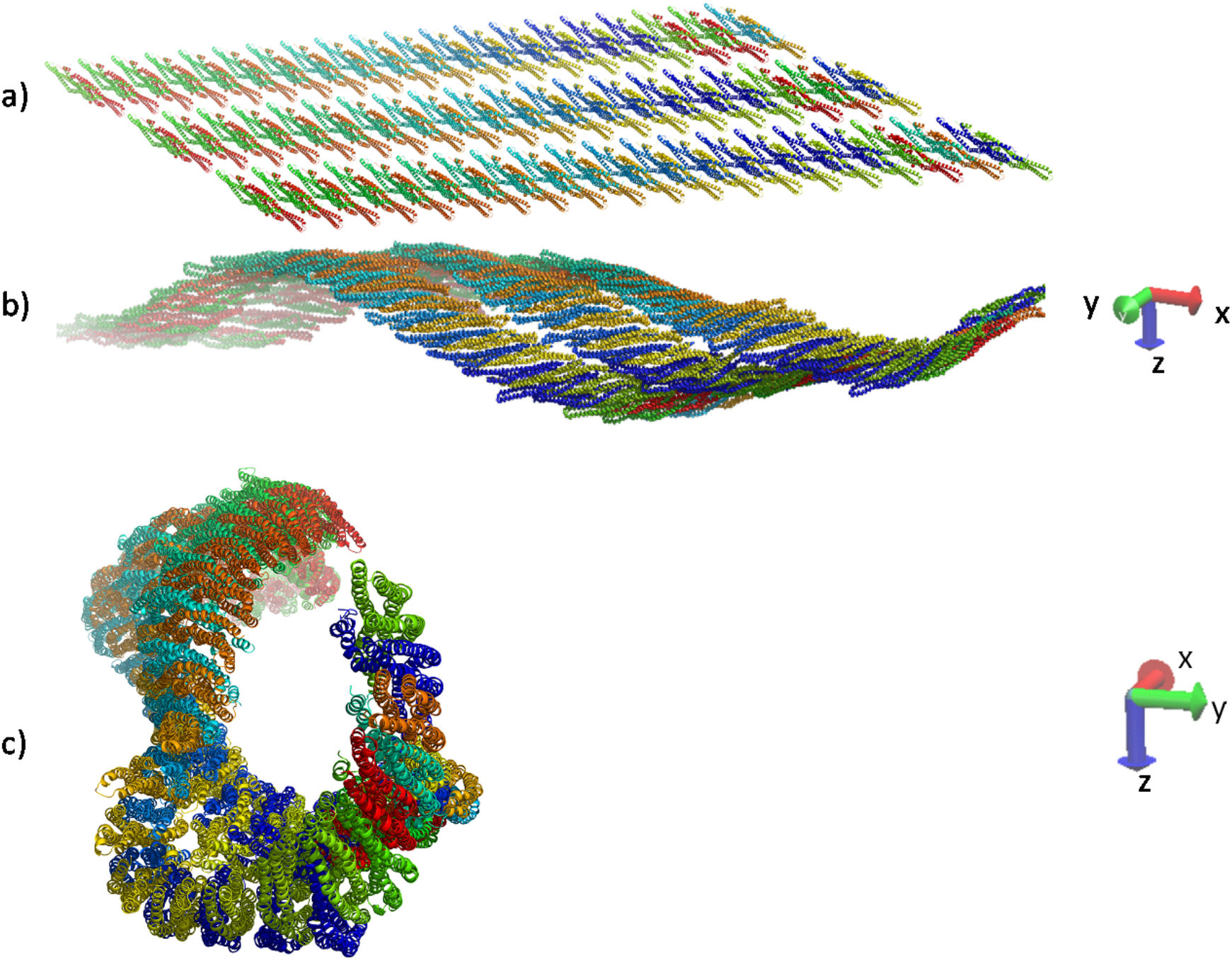
Initial structure of the planar I-BAR dimer sheet (a) and final structure after simulation in implicit water, side view (b) and view along cylindrical axis (c).

As a control, sheets of MIM I-BAR (PDB ID 2D1L) and I-BARa (PDB ID 4NQI) were also simulated in the aqueous phase. Both curved in such a way that the membrane binding surface lay on their convex surface. Interestingly, the curvature developed was smaller than that of IRSP53 I-BAR, consistent with the experimental finding that MIM I-BAR generates larger size tubules than the IRSP53 I-BAR ^14^. Using the same protocol, an F-BAR sheet in aqueous solution (PDB ID 2V0O) developed a curvature with the membrane binding interface on its concave surface, consistent with positive curvature generating behavior. The structures are shown in Fig. S4-S6.

### Simulations of preformed spirals inside a tube

One of the important unresolved questions for I-BAR domains is how they arrange themselves inside membrane tubes. If they behave like other BAR domains, they should also form helical oligomeric structures. It is very difficult to obtain such structures from free simulations of multiple BAR dimers in a reasonable time. Thus, we constructed two types of spirals, with lateral interaction and end-to-end interaction, using information from the free simulations. The helical spiral consisting of lateral interactions is highly stable inside membrane tubes with 20 nm and 40 nm radii (Fig. S7 and S8). The spiral with only end-to-end interactions appeared highly flexible and did not have a definite shape inside the membrane tube (Fig. S9). We also constructed spirals with a combination of lateral and end-to-end interactions where the two helical filaments with end-to-end interactions are laterally overlapping. To design such spirals, the octameric unit (Fig. S10) that was obtained from the free membrane simulations was replicated. The designed spirals with radii 20 nm and 40 nm (Fig. S11) appear rigid and stable throughout the simulations. In both, only the pitch of the helical structure changed to match its curvature preference. In the 40-nm tube the spiral continuously decreases its pitch indicating it prefers a higher curvature. In this oligomeric structure, the orientational preference of individual dimers is satisfied, as they tend to remain parallel to the tube axis, especially inside the 20 nm tube.

Next, linear filaments were constructed with end-to-end interaction between the individual dimers (Fig. S12). The filaments were simulated on the flat membrane surface as well as on the interior of the membrane tube. In both simulations, the filaments failed to give a curved shape. Based on these results we propose that the lateral interaction between the dimers is responsible for negative curvature generation and stabilization.

### Binding energetics of low and high oligomers

The curvature sensitivity of a dimer (Fig. 2c) and a tetramer (Fig. S10) of dimers was examined. The simulations were carried out on a 50% anionic interior tubular membrane. The binding energy (Table 2) is maximal at 10-20 nm and minimal at the flat membrane surface. The curvature sensitivity parameter α obtained from the 20 nm to 50 nm data is 4.0, almost four times that for a single dimer on the same membrane. This suggests that the dimer of dimers tends to have a curved shape and is rigid enough to enhance curvature sensitivity.

**Table 2.** Binding energy (kcal/mol) per dimer in a dimer of dimers and filamentous I-BAR on a 50% anionic membrane. Averages were calculated from three trials and the last 10 ns of a 20-ns simulation.

| Radius | 10nm | 20nm | 30nm | 40nm | 50nm | Flat | $\alpha$ | Sorting Ratio (S) |
| --- | --- | --- | --- | --- | --- | --- | --- | --- |
| Dimer of dimers | -20.6±0.3 | -20.5±0.5 | -19.5±0.6 | -18.9±0.3 | -18.3±0.9 | -18.0±0.6 | 4.0±0.1 | 79.8 |
| Filamentous<sup>a</sup> | | -11.3±0.2 | | -10.2±0.2 | | -9.4±0.2 | 1.63 | 25.0 |

<sup>a</sup> The filaments ( Figure 7, Figure 5, Figure S7) contain 20 dimers.

The binding energetics of the filaments (constructed with lateral interactions) on the flat membrane and the interior of cylindrical membranes with anionic fraction 50% are also presented in Table 2. Binding energies are most favorable for the cylindrical spiral with radius 20 nm and least favorable for the filament on the flat membrane. This is consistent with the generation of smaller-size tubules. The sorting ratio between the flat and 20-nm cylindrical membrane, calculated from the average binding energy of the dimer to the membrane, is ∼25.

## Discussion

The main finding of this work is that oligomers of I-BAR domains attain a curvature that is unrelated, in fact nearly orthogonal, to the intrinsic curvature of the dimer along its principal axis. The curved oligomer exposes a convex membrane binding surface and is thus consistent with generating negative membrane curvature. Both lateral and end-to-end interactions contribute to oligomerization, but the former are stronger. The relevant curvature develops along these lateral interactions. As a control, an F-BAR domain oligomer develops curvature in the opposite direction, consistent with positive membrane curvature generation. Binding energy calculations show that oligomerization increases the curvature sensitivity of the I-BAR domains.

Experimental evidence on I-BAR oligomerization is scarce. Slow fluorescence recovery in the absence of stiffening of nanotubes coated with high density of ABBA I-BARs indicated possible oligomerization but with flexible linkages ^27^. Similar results were obtained with other BAR domains ^28^. Only a slight tendency for oligomerization of I-BAR domains has been observed in the aqueous phase ^19^. The fact that our oligomers are stable in water may be due to the brief duration of the simulations or, more likely, to an overestimation of the driving forces for protein adhesion by the implicit solvent model. However, even if the stability of the oligomers is overestimated, the qualitative trends regarding curvature should be valid.

What causes the different curving of I-BAR and F-BAR oligomers? Some F-BARs, such as FBP17 and CIP4 ^29^ have enough intrinsic curvature as dimers to generate tubules by orienting perpendicular to the tube axis and the radius of the tubules they generate matches their intrinsic curvature. For these, oligomerization is needed only to provide sufficient protein density. Others, such as Fcho2 ^30^ and the pacsins ^31,32^, have kinked “wings” that give them an S-shape when viewed from above, while I-BARs are straight. The binding interface between dimers is very different for the BAR domains we considered here (Fig. 5). I-BAR dimers are parallel to each other with a lateral shift; the contact interface is extensive and the salt bridges that drive oligomerization occur mostly in helix 3, which is the one farthest from the membrane. This likely causes the membrane binding surface to curve in a convex way. The Fcho2 F-BAR appears to oligomerize end-to-middle and the wings make salt bridges in helix 1, close to the membrane, which likely causes the membrane binding surface to curve in a concave way. These structures provide many opportunities for validation by mutagenesis or distance measurements via DEER or FRET.

**Figure 5.**
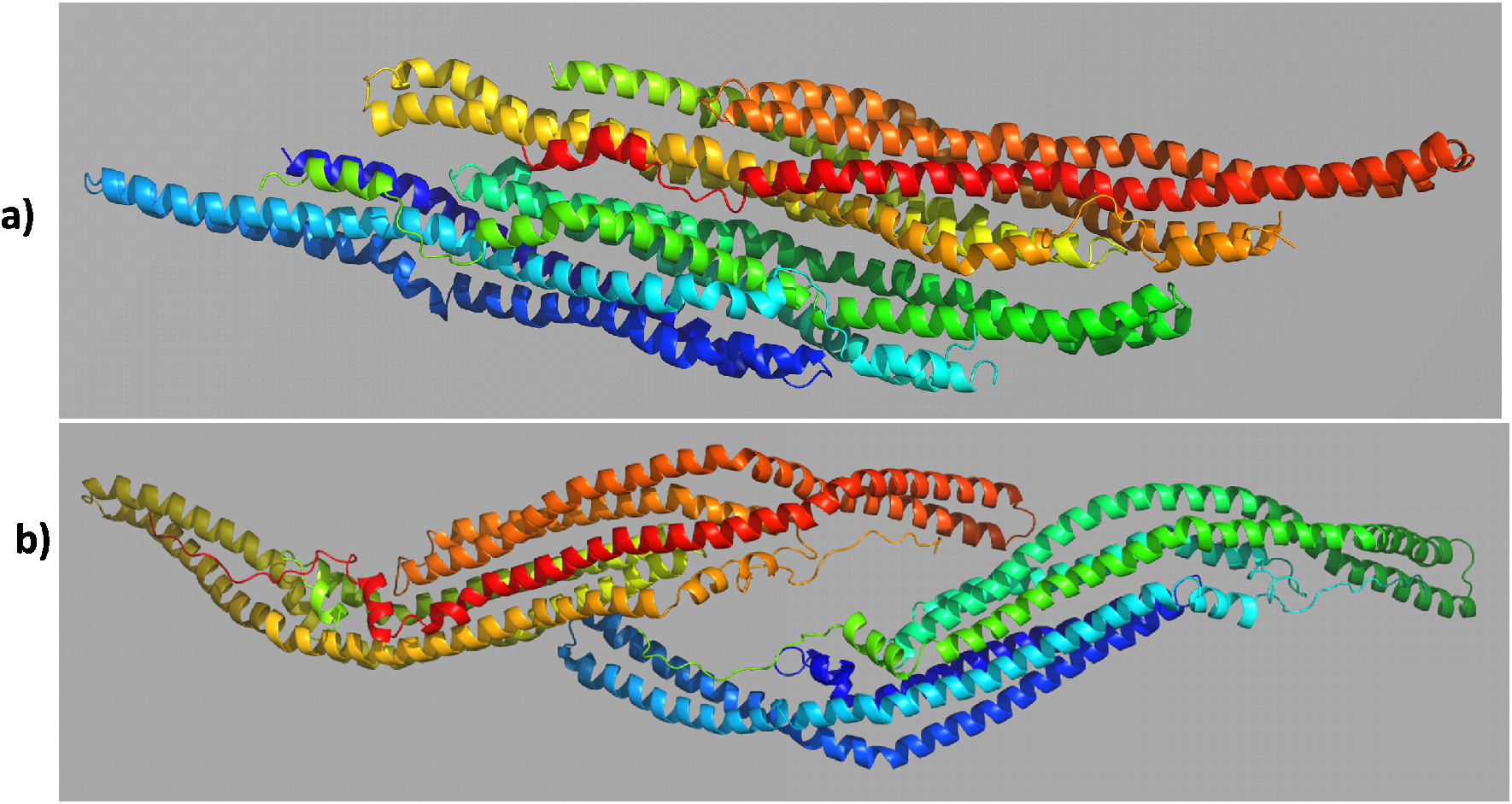
Dimer of dimers of I-BAR (a) and F-BAR (b) viewed from the top as they interact in the planar sheets of Fig. 4 and S5, respectively.

Strictly speaking, since our implicit membranes are rigid, what we have measured is curvature sensing, not curvature generation. However, these processes are thermodynamically linked. A curvature sensor will generate curvature if the protein density and the differential membrane binding energy are sufficient to overcome the cost of membrane deformation. We can verify this using our results together with elasticity theory^33^. Consider a membrane of area A and N I-BAR dimers. We bend the membrane into tubes of radius R and length L (A=2πRL). For large enough A we can neglect edge effects. If the percent surface coverage is φ and the area per dimer is α, N/A = φ/α. The binding energy per dimer for oligomeric I-BAR relative to the flat membrane (Table 2) can be modelled as E_b_ = −40/R kcal/mol down to R ∼ 20 nm and is most favorable at 20 nm. The energy of the tube relative to the flat membrane is E = πκL/R + N E_b_, or E/A = κ/2R^2^ – φ/α 40/R, where κ is the bending rigidity. The equilibrium radius is obtained by setting dE/dR = 0, giving R_eq_=κα/40φ. For κ=12.5 kT^19^, α=50 nm^2^, φ=0.25, and T=298 K we obtain R_eq_ = 37.5 nm. This is within the range of tubules observed experimentally^14,34^. The balance between membrane deformation energy and protein binding leads to radii larger than the optimal 18 nm observed in preformed tubule experiments^19^ and the 13 nm curvature that the IRSp53 I-BAR oligomer develops spontaneously in our implicit water simulation. Clearly, the differential binding energies we calculate per dimer in the oligomeric state are sufficient for membrane curvature generation at reasonable surface densities.

In pulled tube experiments optimal partitioning is observed for 18-nm tubes ^19^. Our binding energy calculations reproduce the stronger binding upon increasing the curvature to 20 nm and the reduction in binding from 20 nm to 10 nm at higher anionic fractions. At 30% anionic fraction, the binding affinity is about the same at 10 and 20 nm radius. The pulled tube experiments ^19^ and another study ^27^ found that the sorting ratio (partition coefficient) is higher at lower protein densities. This seems consistent with the formation of specific oligomeric structures in the tubule interiors with a definite, optimal protein density. As protein concentration increases, less optimal structures and curvatures are populated on the flat membrane and the sorting ratio decreases. The experiments showed that the sorting ratio was 20, 12 and 5 at overall protein area fraction 1%, 2%, and 5%, respectively. At these conditions, the surface coverage on the pulled tubes was 20, 24 and 25%, respectively. The surface area coverage of our spiral on Fig. S7 is ∼ 8.2%, about 3 times lower, but it is feasible to construct tighter spirals with higher surface coverage. The sorting ratio calculated from our binding free energies of the individual dimer is 1.8, 2.7 and 6.4 in 30%, 50% and 75% anionic membranes, respectively (Table S3). For the oligomeric forms the sorting ratio is substantially higher. For example, for the 20-dimer oligomeric spiral in 50% anionic membrane it is ∼25 (Table 2). Precise determination of the sorting ratio requires detailed consideration of oligomerization equilibria and determination of accurate oligomeric structures on both the flat membrane and the tube.

Several studies suggested that the lipid PI(4,5)P2 (PIP2) plays a role in I-BAR function ^14,34^, although it is not strictly required for membrane remodeling ^27,35^, suggesting no specific binding to BAR domains. At neutral pH PIP2 has a charge of −4 ^36^ and its clustering by I-BAR ^14^ further increases the effective charge of the membrane and thus the binding energy. Our implicit membrane model neglects specific lipid binding effects and assumes uniform distribution of charge. Thus, to reproduce the behavior of a PIP2-containing membrane we need to use higher membrane anionic fractions than the nominal value. Considering this, the 50 or 70% anionic fractions where we observe the greatest curvature sensitivity are not unreasonable. In addition, Folch fraction I lipids commonly used in *in vitro* experiments ^31^ have similar anionic fractions. ^37^

Previous MD simulations found that the IRSp53 domain flattens on the membrane and appears too flexible to support membrane remodeling based on its own intrinsic curvature ^22^. This is consistent with our results. Another all-atom simulation with a single I-BAR dimer could not generate the expected high negative curvature, implying that some sort of oligomerization must play a role ^21^. Jarin et al. ^23^ used two levels of coarse-graining to study the aggregation behavior of I-BAR dimers on membranes of various geometries. They found that I-BAR aggregates oriented parallel to the tube axis when the tube radius was narrow (25 nm) and perpendicular when the tube was wider (radius 50 nm). This is in agreement with our results. Another highly coarse-grained model explored the effect of PIP2 clustering and found that preformed protrusions were stabilized by I-BAR dimers ^24^. Consistent with our results, they mainly observed two types of interactions; side-by-side and end-to-end, and these interactions were controlled by the strength of the membrane-protein interaction, protein curvature and the inclusion of the PIP2 patches on the membrane. However, these models employed purely repulsive protein-protein interactions and focused on membrane-mediated interactions. Thus, they could not predict oligomers of definite shape. Our results highlight the importance of protein-protein interactions and could be used to develop more realistic coarse-grained models that capture such effects and allow the study of detailed pathways of membrane deformation.

## Methods

### IMM1_curv Model

Implicit membrane modeling was carried out with IMM1_curv ^26^, which can model spherical or cylindrical membranes of specified radius. IMM1 ^38^ is an implicit membrane model based on the effective energy function 1 (EEF1) ^39^. EEF1 is based on the polar-hydrogen-only CHARMM force field and a solvation free energy which is calculated as the sum of group contributions:

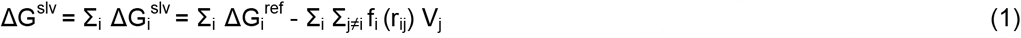

where ΔG_i_^slv^ is the solvation free energy of atom i and ΔG_i_^ref^ is the solvation free energy of atom i in a small model compound. The last term describes the loss of solvation due to surrounding groups; f_i_ is the solvation free energy density of atom I calculated as a Gaussian function of r_ij_, r_ij_ is the distance between atoms i and j, and V_j_ is the volume of atom j. Interactions between the proteins, critical in this work, are governed by the common interatomic Lennard-Jones and Coulomb potentials, in addition to a desolvation term.

In IMM1, ΔG_i_^ref^ is modified to account for heterogenous membrane-water systems.

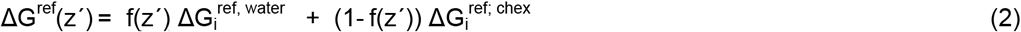

where ΔG_i_^ref,water^ and ΔG_i_^ref;chex^ are reference solvation free energies of atom i in water and cyclohexane, respectively. f(z’) is a sigmoidal function that describes the vertical transition from one phase to the other:

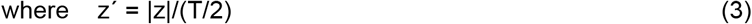

where T is the thickness of the hydrophobic core of the membrane. The exponent n controls the steepness of the transition. The increase in electrostatic interactions inside a membrane is taken into account by a modified dielectric screening function given by,

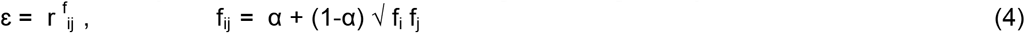

where α is an empirical parameter and was adjusted to 0.85 which gives membrane binding energies close to experiment.

IMM1_curv modifies the shape of the hydrophobic region to account for a cylindrical or spherical shape. The center of the sphere is placed at the origin and R is the distance between the origin and the sphere mid-surface. The equation for z above is modified to z’ = |r-R|/(T/2) where r is the distance of the atom from the origin, given by

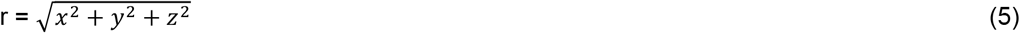

For a cylinder the x component is omitted, assuming that the cylindrical axis is along the x axis. The lateral pressure profile further takes account the lipid packing effect as a result of the curvature. Approximate Poisson-Boltzmann equations were used to calculate the electrostatic interaction between the protein and anionic membrane with spherical and cylindrical shapes.

### Computational Details

The MD simulations were carried out with the CHARMM package ^40^. The simulations were run for 20 ns at 300 K with a 2-fs time step. Due to the lack of solvent friction, the true time-scale of implicit solvent simulations is much longer than their nominal duration. As a result, this duration was sufficient for achieving structural and energetic convergence (see Figs. S13-S20 in SI). The trajectories were saved every 1 ps and analysis was done on the last 10 ns. The initial structure of the dimer was obtained from PDB code 1Y2O ^41^. In the IMM1 calculations, the width of the membrane hydrophobic core was set to 25.4 Å and the anionic fraction was set to 30%. For the higher oligomers, 50% anionic membrane was used. The charge layer was set 3 Å outward from the hydrophobic-hydrophilic interface and 0.1 M salt concentration was used. Since membrane binding is dominated by electrostatic interactions, calculations were carried out without the lateral pressure component. Multiple trials were averaged to estimate statistical uncertainty.

The binding energies were estimated as average transfer energies, i.e. the difference in energy of the peptides on the membrane surface and the same conformation in bulk water. This is approximate, because it neglects possible changes in intramolecular energy (including such changes makes the results very noisy). Conformational and translational entropy is also neglected in these calculations, but these contributions are likely to be very similar in binding to flat and curved membranes. Two-sample t-tests were done to verify the statistical significance of binding energy differences for different curvatures (Table S4).

A quantitative measure of curvature sensitivity is the parameter α defined by the equation below ^26^.

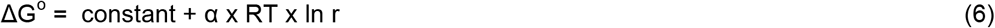

where r is the radius of the curvature.

## Acknowledgments

This work was supported by the NIH (GM117146) and the NSF (MCB 1855942). Infrastructure support was provided in part by Research Centers in Minority Institutions grant no. 8G12MD007603 from the NIH.

## Author Contributions

BN performed the bulk of the computations. AS contributed in the construction of the oligomers. BN and TL designed the research and wrote the manuscript.

