## Supplemental Information for "Mechanism of negative membrane curvature generation by I-BAR domains"

**Table S1**. Electrostatic contribution of positively charged residues to binding on a 30% anionic membrane surface. The first value is for a flat membrane and the second value inside a spherical membrane of 30 nm radius. The averages are calculated from the last 2 ns of the 20-ns simulation. The energies are in kcal/mol per monomer.

| | | | |
|---|---|---|---|
| LYS 18 | -0.41, -0.33 | LYS 147 | -0.70, -0.96 |
| LYS 36 | -0.41, -0.25 | LYS 152 | -0.24, -0.42 |
| LYS 40 | -0.45, -0.27 | LYS 156 | -0.39, -0.66 |
| LYS 121 | -0.80, -0.53 | ARG11 | -0.16, -0.38 |
| LYS 122 | -0.48, -0.25 | ARG 29 | -0.47, -0.28 |
| LYS 136 | -0.74, -0.70 | ARG 114 | -0.51, -0.35 |
| LYS 142 | -0.29, -0.40 | ARG 128 | -0.54, -0.57 |
| LYS 143 | -0.73, -0.90 | | |
| LYS 146 | -0.44, -0.70 | | |

**Table S2.** Tilt angle of an I-BAR dimer with respect to the tube axis. Initially, the protein was placed parallel to the tube axis except one trial, which was started in a perpendicular orientation.

| | 20nm | 30nm | 40nm | 50nm |
|---|---|---|---|---|
| Inside | | | | |
| Trial 1 | 12±3 | 19±3 | 85±9 | 3±5 |
| Trial 2 | 8±4 | 29±2 | 6±2 | 90±3 |
| Trial 3 | 6±3 | 17±2 | 23±2 | 84±2 |
| Trial 4 | 17±3 | 3±2 | 29±2 | 7±3 |
| Perpendicular | 16±9 | 30±3 | 88±3 | 85±6 |
| outside | | | | |
| Trial 1 | 12±2 | 10±2 | 10±4 | 13±2 |
| Trial 2 | 6±2 | 14±1 | 21±3 | 13±2 |
| Trial 3 | 13±2 | 1±2 | 2±2 | 10±2 |
| Perpendicular | 20±3 | 28±2 | 15±3 | 12±3 |

**Table S3**. Sorting ratio between the planar and tubular membrane (radius 20 nm) at various anionic fractions.

$S = \exp(-\{\Delta G_{20nm} - \Delta G_{plane}\})\ /K_bT)$

| System | Sorting Ratio (S) |
|---|---|
| 30 % anionic | 1.82 |
| 50% anionic | 2.71 |
| 75% anionic | 6.36 |

**Table S4**. Student's t-test analysis of differences in binding energies between the cylinder (20 nm, inside) and the flat membrane (Tables 1 and 2). The difference is statistically significant when it is larger than the uncertainty. Energies are in kcal/mol. DF is the number of degrees of freedom.

| system | Confidence Level | | | t | P-values | DF (N1+N2-2) |
|---|---|---|---|---|---|---|
| | 95% | 90% | 80% | | | |
| Dimer (30% anionic) | -0.3±0.6 | -0.3±0.5 | -0.3±0.4 | -0.94 | 0.38 | 6 |
| Dimer (50% anionic) | -0.6±0.4 | -0.6±0.3 | -0.6±0.3 | -2.91 | 0.05 | 6 |
| Dimer (75% anionic) | -1.1±0.7 | -1.1±0.6 | -1.1±0.4 | -3.11 | 0.02 | 6 |
| Dimer-of-dimers (50% anionic) | -2.5±0.9 | -2.5±0.7 | -2.5±0.6 | -6.76 | 0.006 | 4 |
| Lateral Spiral, per dimer | -1.9±0.04 | -1.9±0.03 | -1.9±0.02 | -212.43 | 0.001 | 1998 |

1a)

1b)

1c)

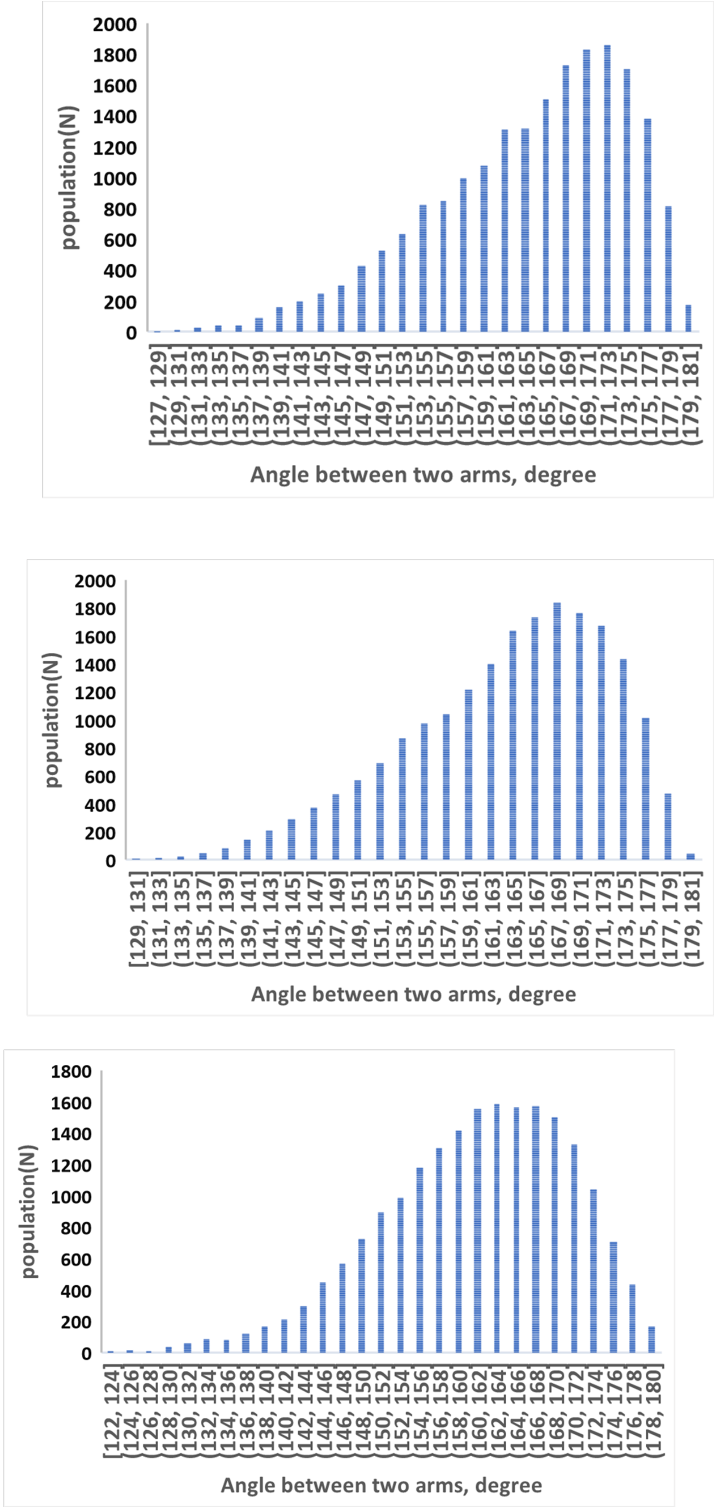


**Figure S1**. Distribution of the angle between two arms during entire 20 ns simulation. The angle is defined in Fig 1c. In the crystal structure it is ~154º. a) flat membrane b) 30 nm spherical surface (outside) c) 30 nm spherical surface (inside)

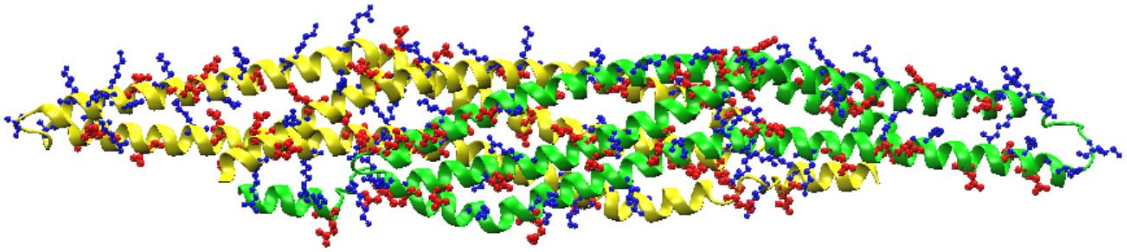

**Figure S2**. Structure of a dimer of dimers with charged residues shown in ball and stick model, ARG/LYS in blue and ASP/GLU in red.

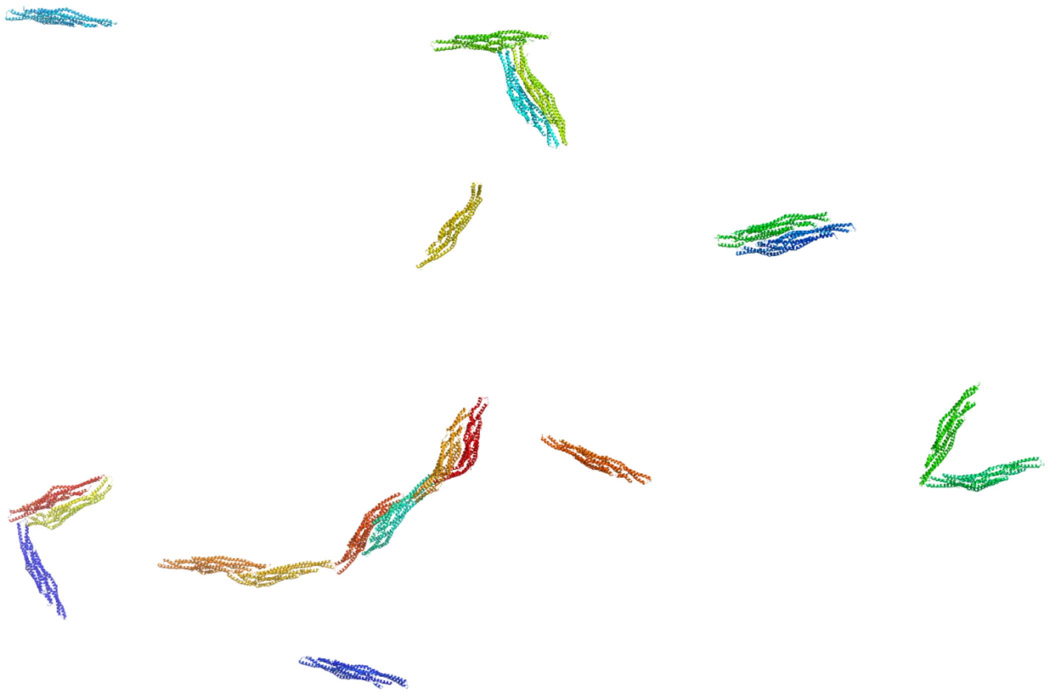

**Figure S3**. Free simulation of 20 dimers on a flat membrane surface.

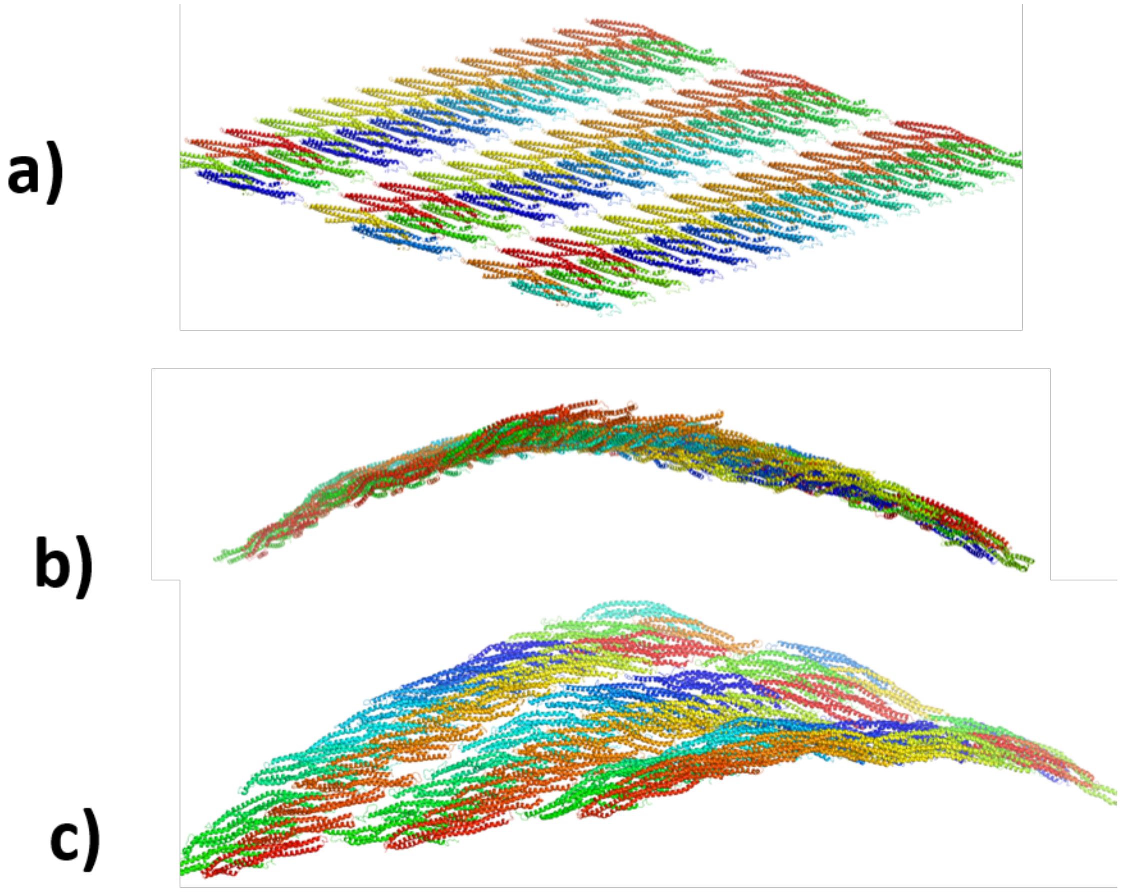


**Figure S4.** Solution MD simulation of MIM-IBAR  planar sheet. a) initial structure, b) side-view after 8 ns simulation,  c) top-view after 8 ns simulation.

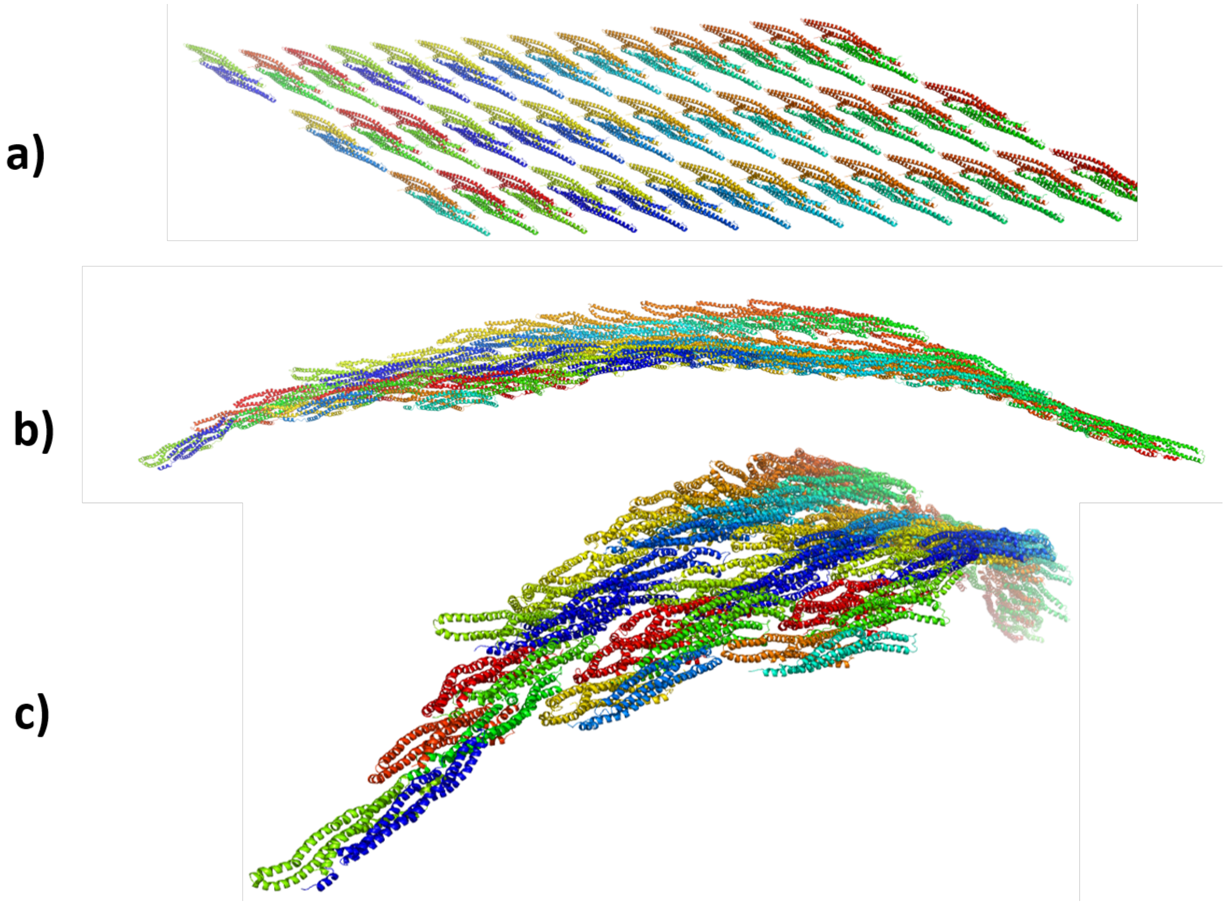


**Figure S5.** Solution MD simulation of IBARa (PDB ID 4NQi) planar sheet. a) initial structure, b) side-view after 8 ns simulation,  c) top-view after 8 ns simulation.

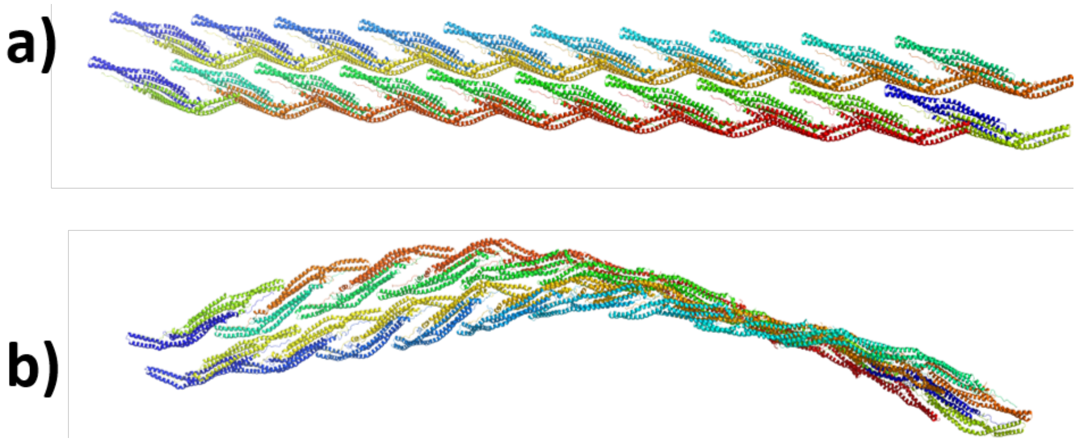


**Figure S6.** Solution MD simulation of F-BAR (PDB ID 2V0O) planar sheet. a) initial structure, b) final structure after 30 ns simulation

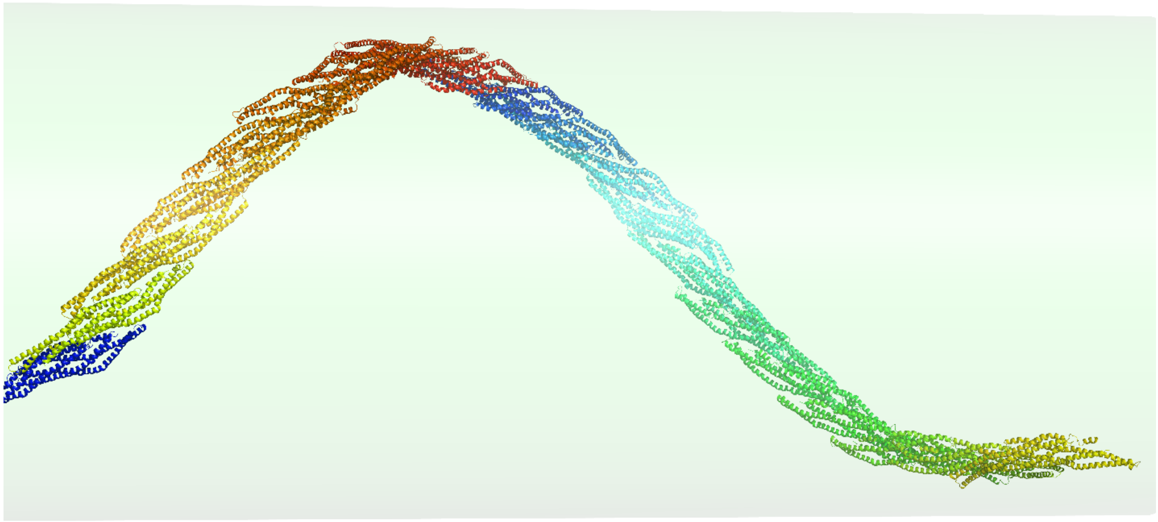

**Figure S7.** Final configuration of the pre-constructed spiral after 20 ns simulation in the interior of a 20-nm cylindrical tube. The initial and final pitch of the helix is 110 nm and 107 nm, respectively.

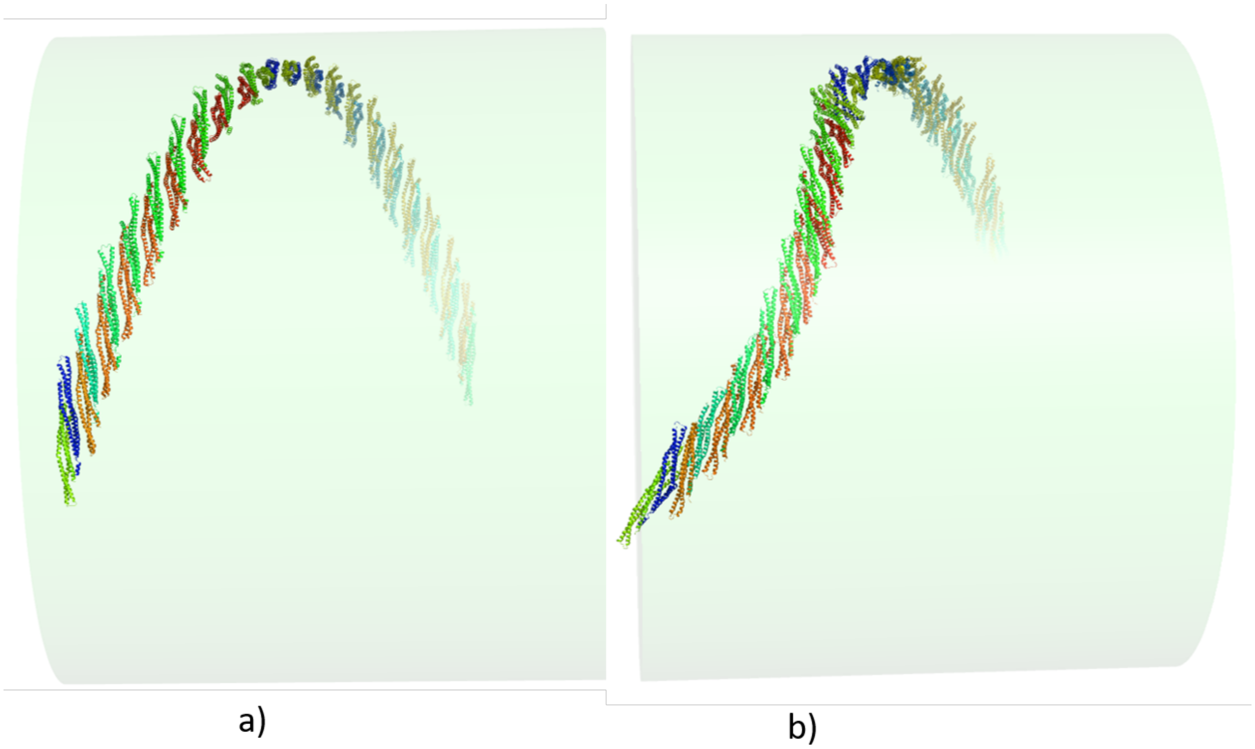


**Figure S8**. The constructed helical spiral inside the tube with radius 40 nm with lateral interactions between the monomers, a) initial b) final after 30 ns simulation

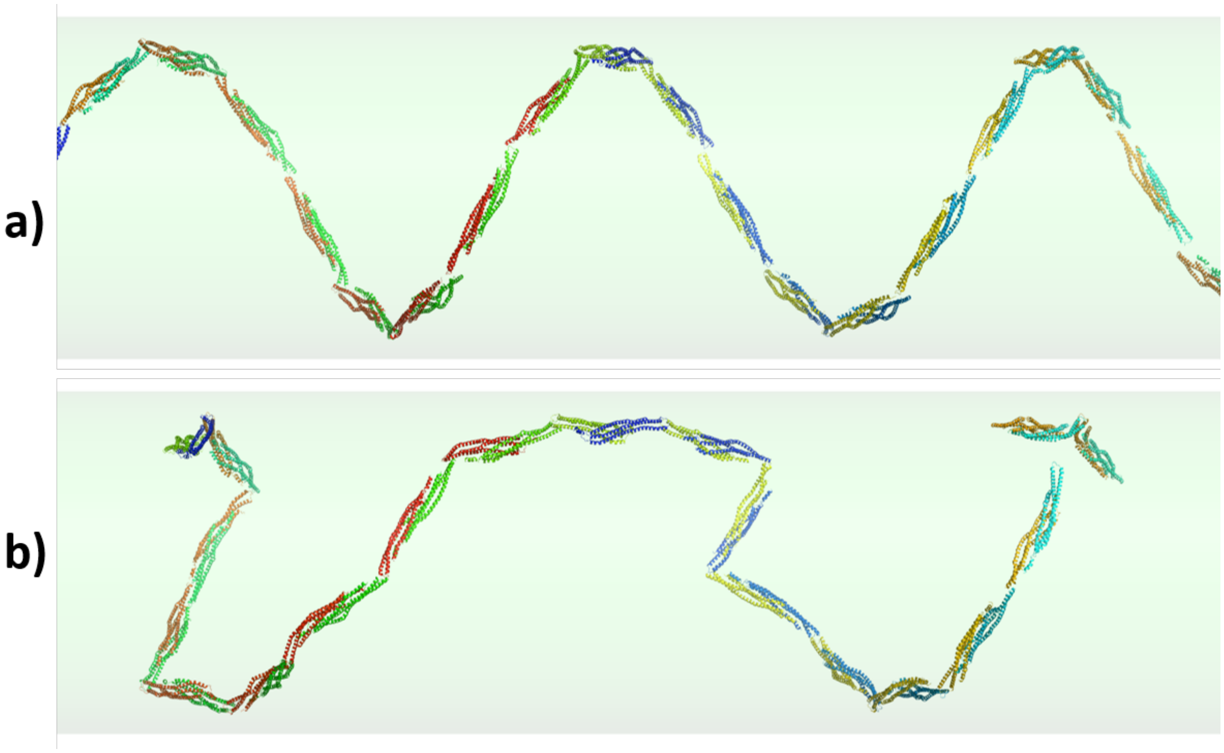


**Figure S9.** Constructed helical spiral with end-to-end interaction with radius 20 nm a) initial and b) after 10 ns simulation.

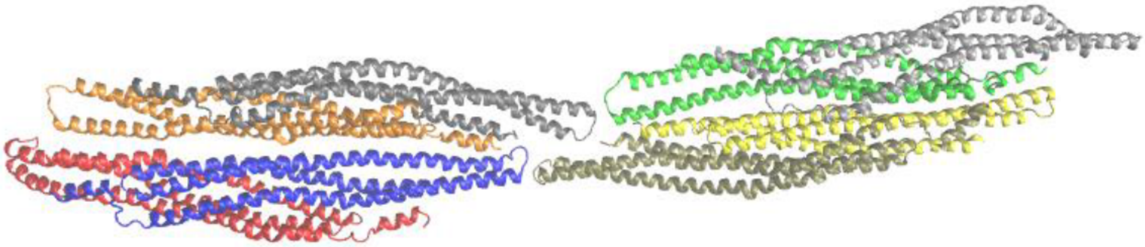

**Figure S10**. Structure of tetramer of dimers obtained from the free simulation of multiple dimers on the flat membrane surface.

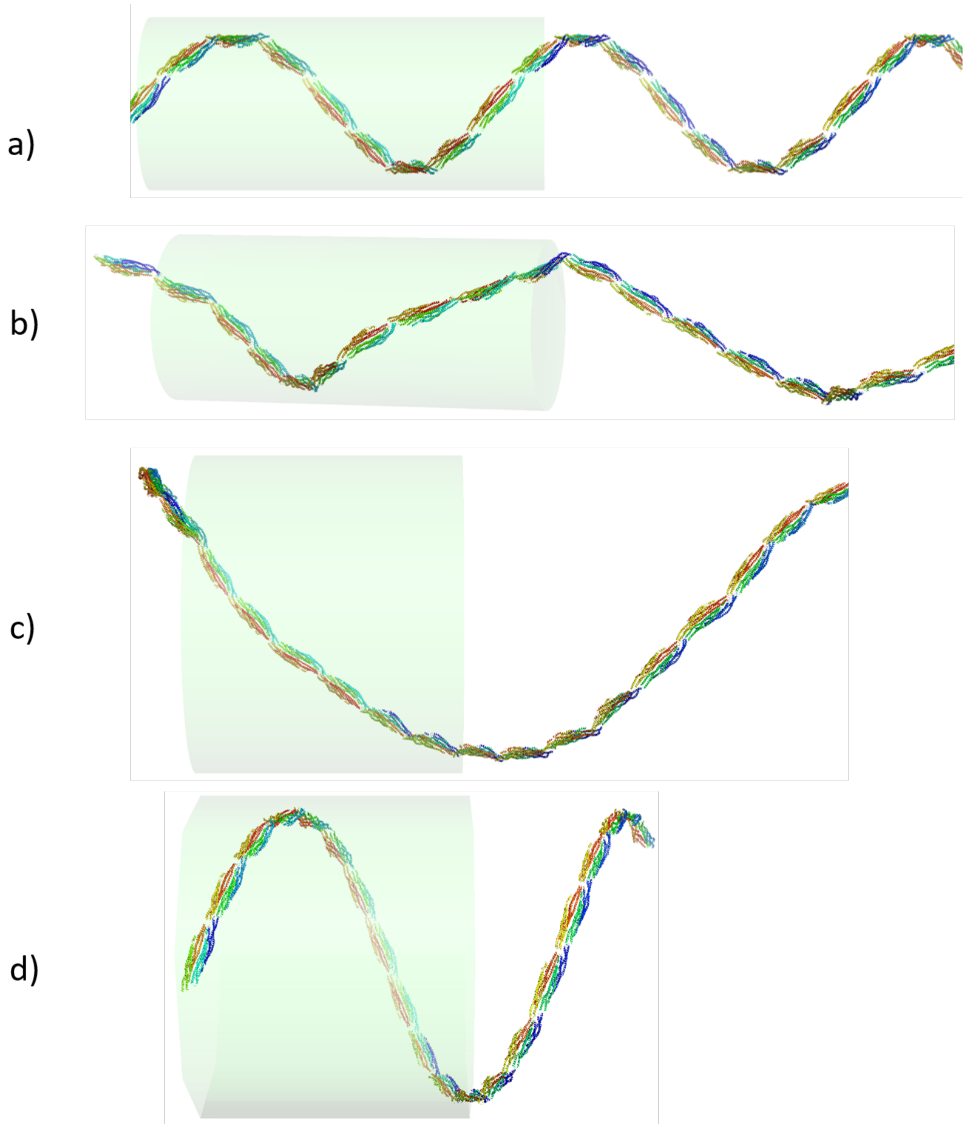


**Figure S11.** Constructed helical spiral inside a tube with radius 20 nm a) initial (helical pitch 86 nm), b) final after 20 ns simulation (helical pitch 134 nm). 40-nm constructed spiral c) initial (helical pitch 168 nm) and d) final after 20 ns simulation (helical pitch 83 nm). The spirals were designed by laterally overlapping filaments with end-to-end interactions.

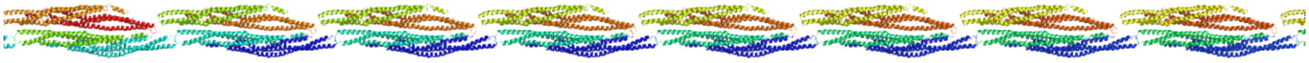

**Figure S12.** Linear filament designed by laterally overlapping two filaments with end-to-end interactions between the dimers.

**Figure S13**. Energy vs. time plot for the I-BAR dimer simulations on 20-50 nm tube interior (30% anionic)

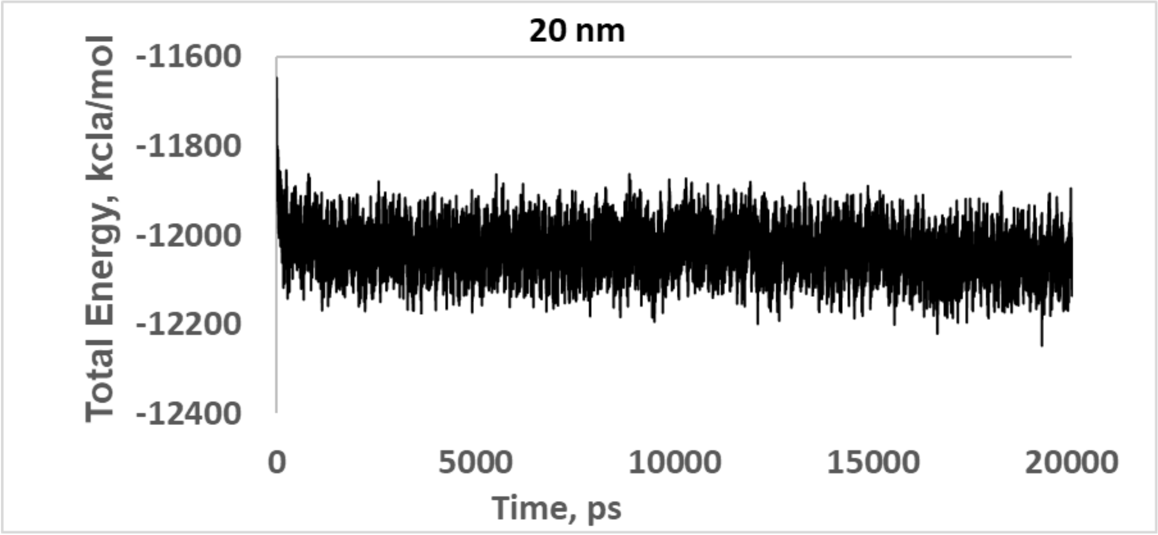

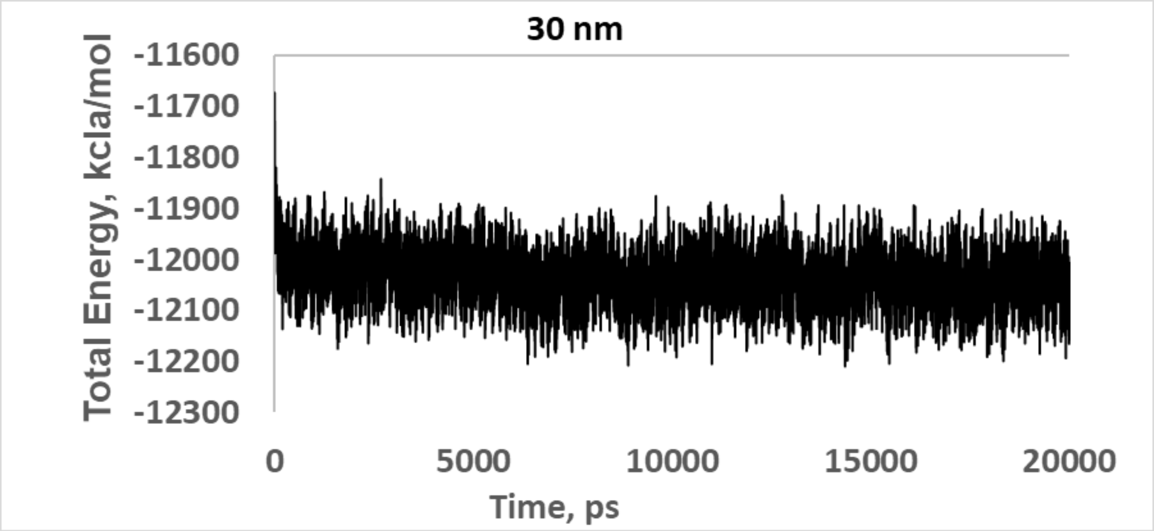
30 nm
Total Energy, kcla/mol
-11600
-11700
-11800
-11900
-12000
-12100
-12200
-12300
0
5000
10000
15000
20000
Time, ps

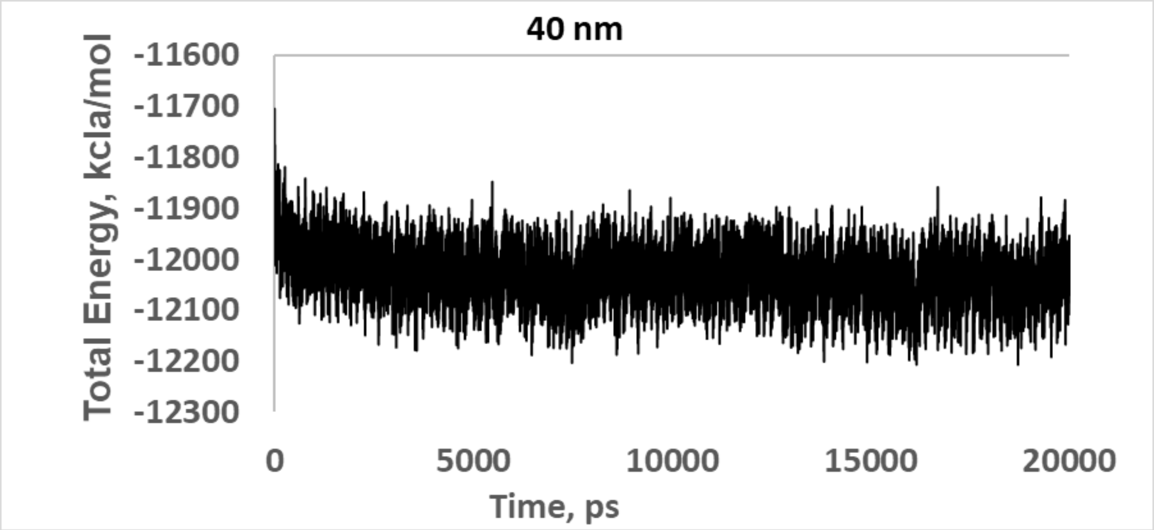
40 nm
Total Energy, kcla/mol
-11600
-11700
-11800
-11900
-12000
-12100
-12200
-12300
0
5000
10000
15000
20000
Time, ps

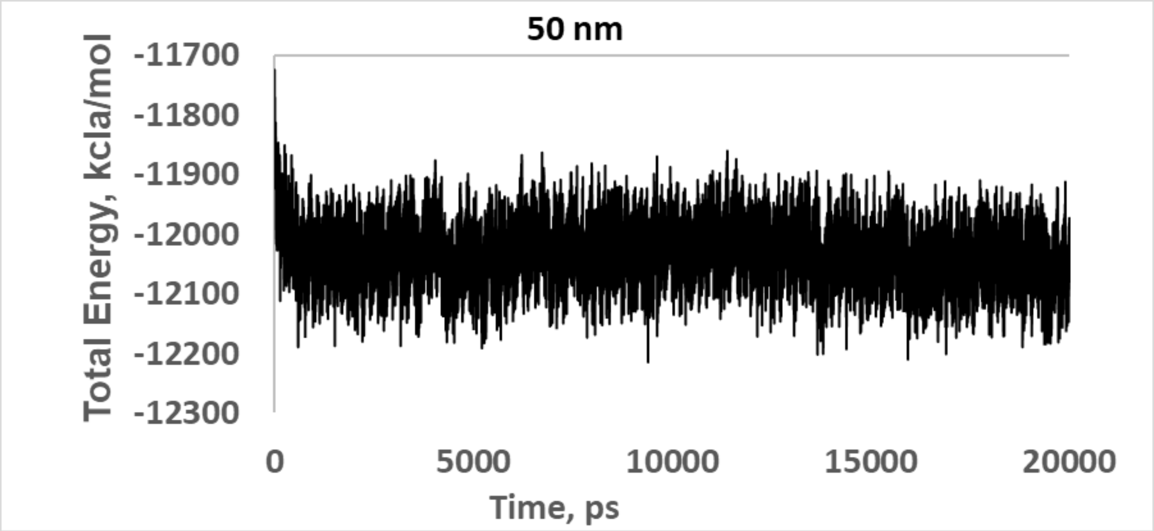
50 nm
Total Energy, kcla/mol
-11700
-11800
-11900
-12000
-12100
-12200
-12300
0
5000
10000
15000
20000
Time, ps

**Figure S14**. Energy vs. time plot for I-BAR dimer-of-dimers inside a 20-50 nm tube for 50% anionic membrane.

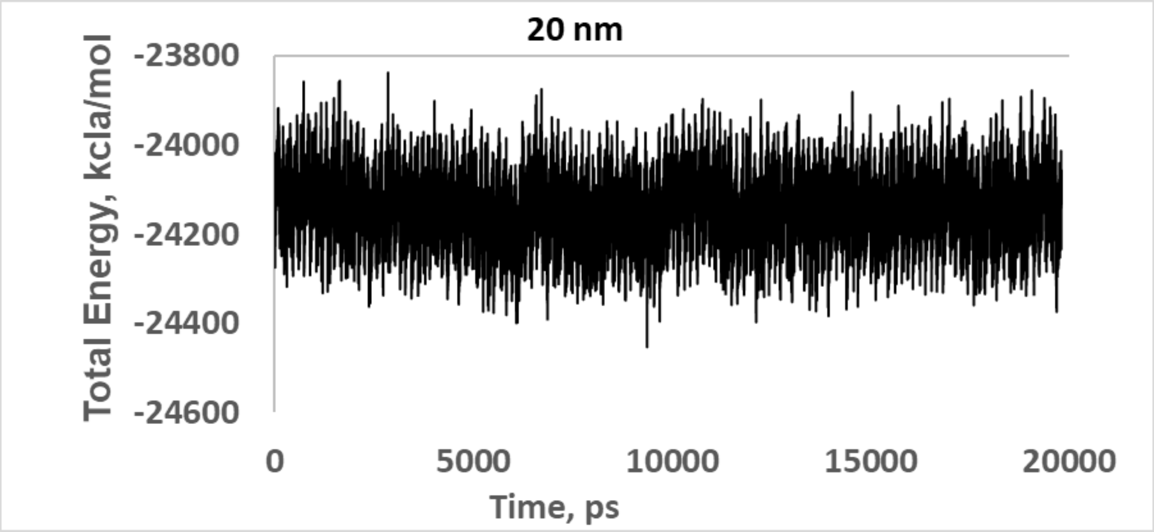


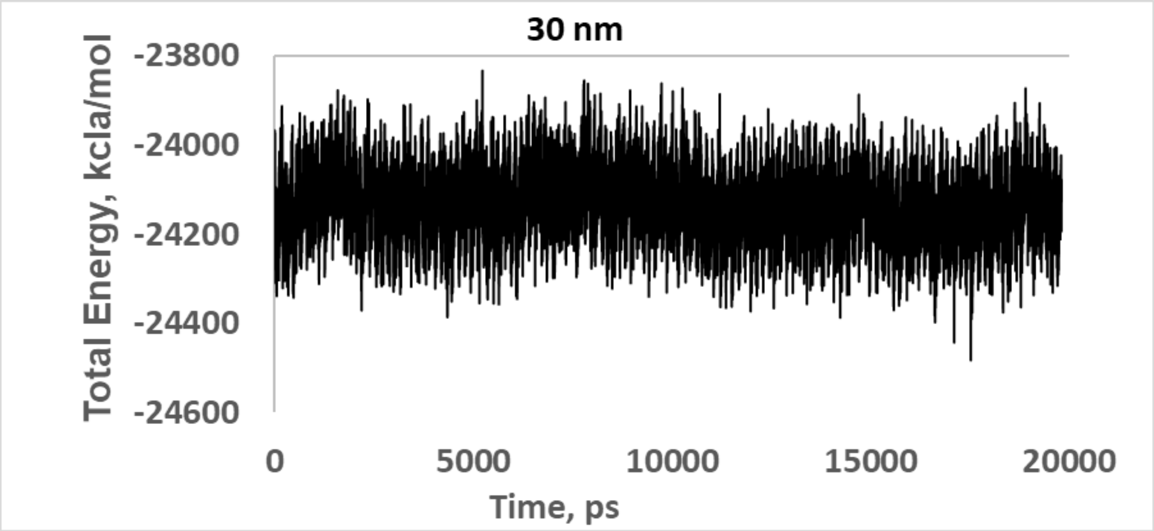

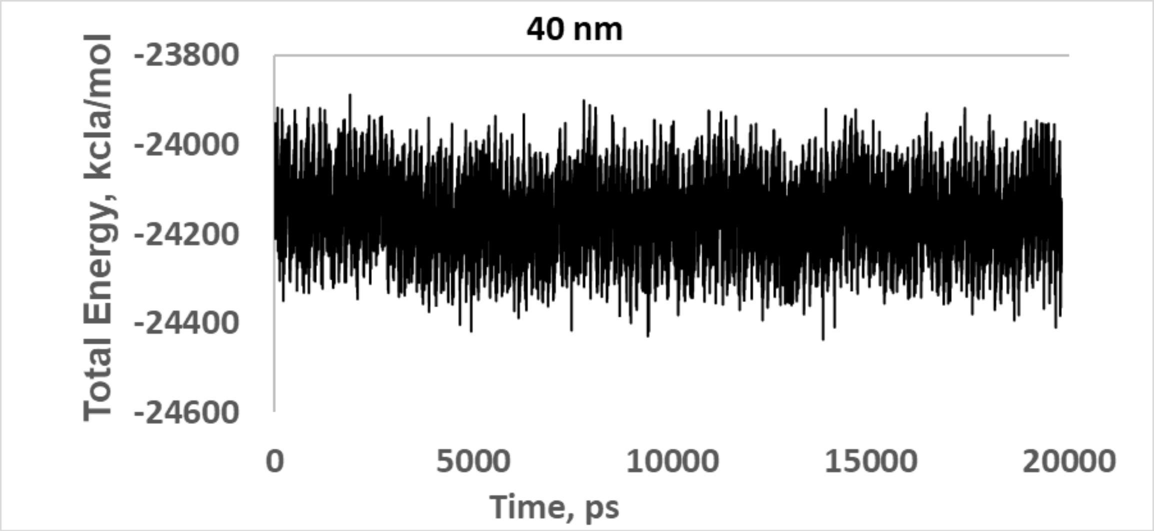
40 nm
Total Energy, kcla/mol
-23800
-24000
-24200
-24400
-24600
0
5000
10000
15000
20000
Time, ps

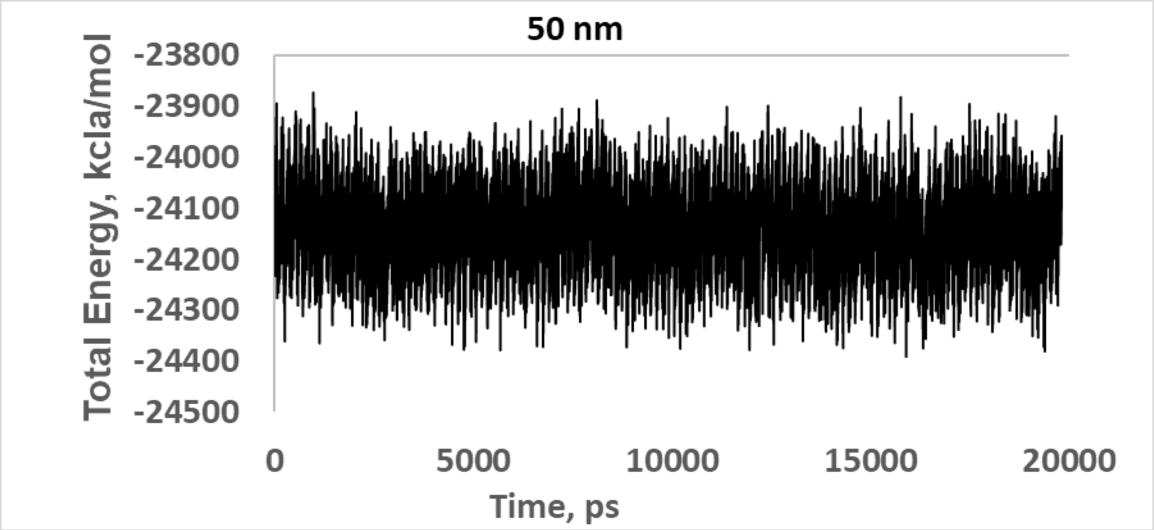
50 nm
Total Energy, kcla/mol
-23800
-23900
-24000
-24100
-24200
-24300
-24400
-24500
0
5000
10000
15000
20000
Time, ps

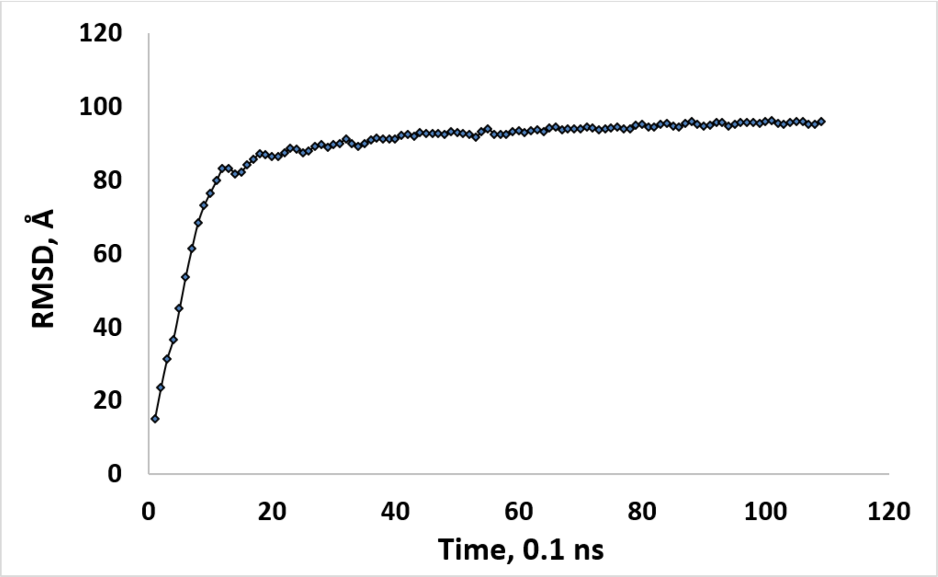


**Figure S15**. RMSD plot for the implicit water simulation of IRSp53 I-BAR sheet

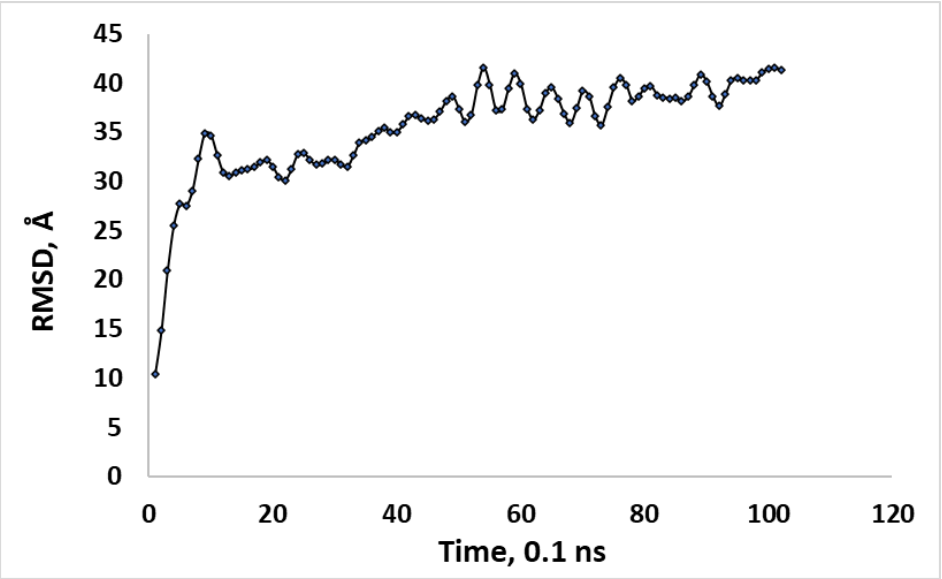


**Figure S16.** RMSD plot for the implicit water simulation of MIM I-BAR sheet (Fig S4)

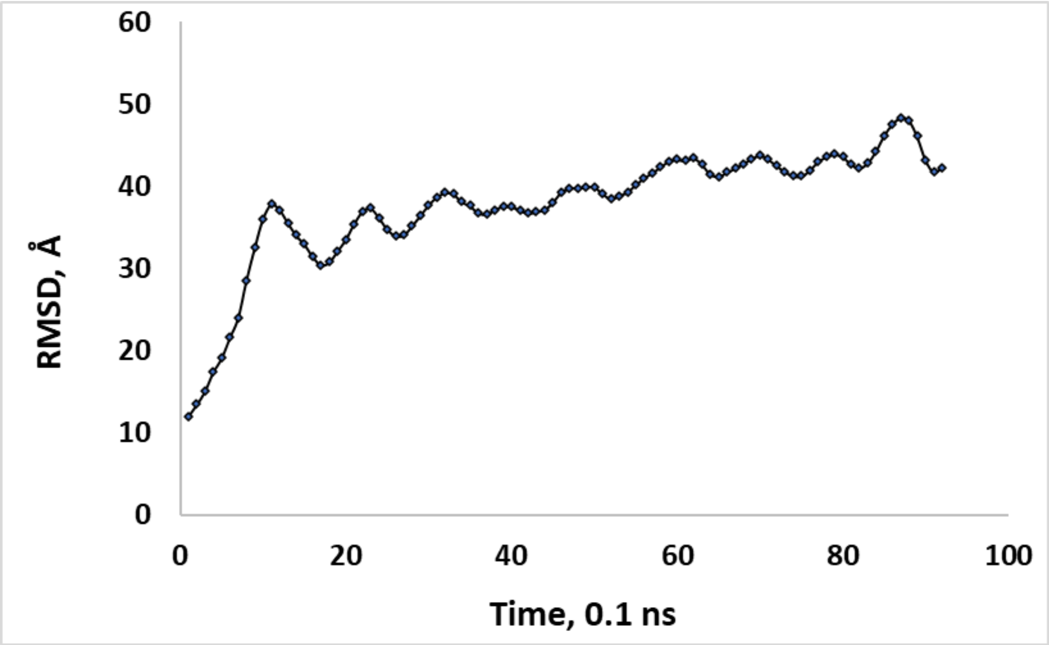


**Figure S17**. RMSD plot for the implicit water simulation of I-BARa sheet (FigS5)

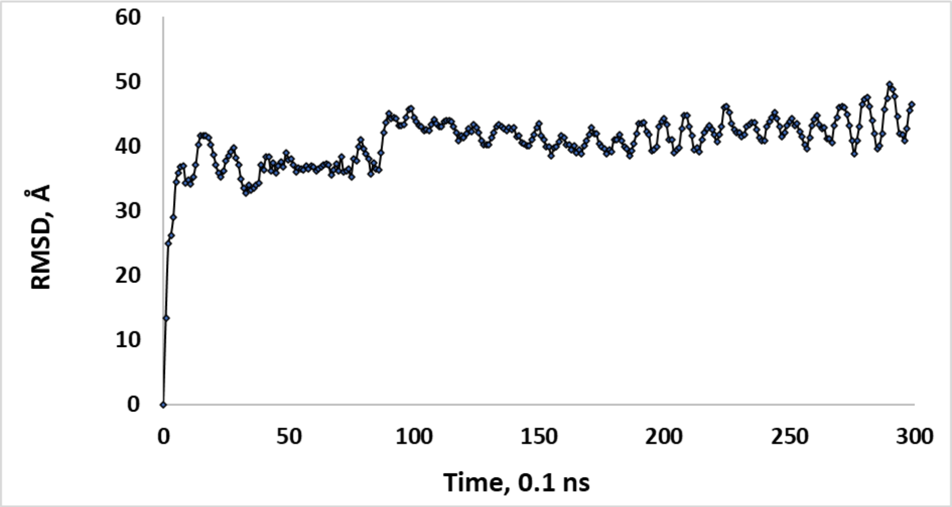


**Figure S18**. RMSD plot for the implicit water simulation of F-BAR sheet (FigS6)

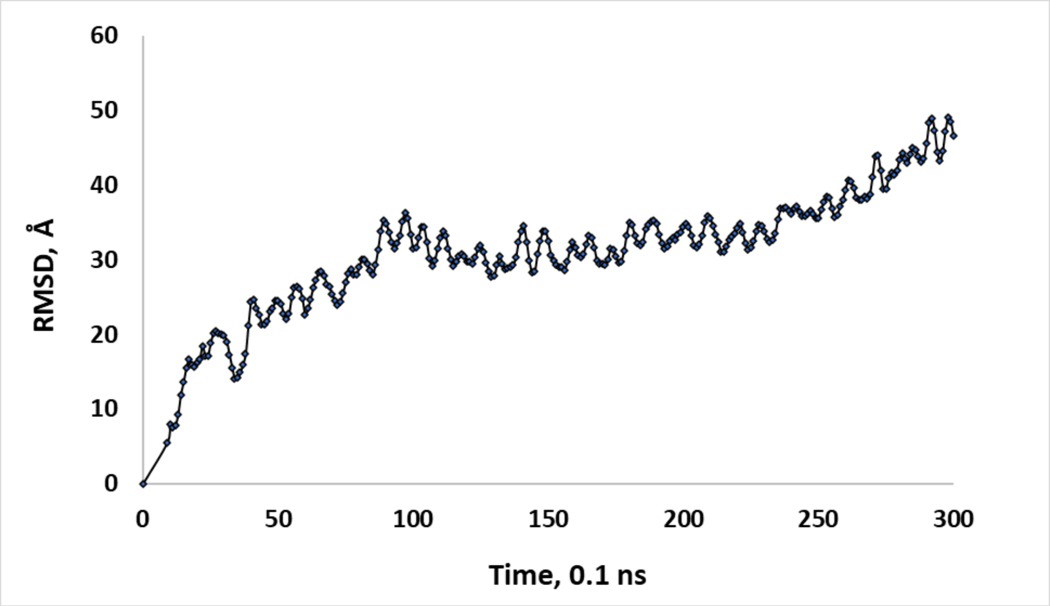


**Figure S19** RMSD plot of the pre-designed spiral with lateral interactions inside the 20 nm tube (Fig S7)

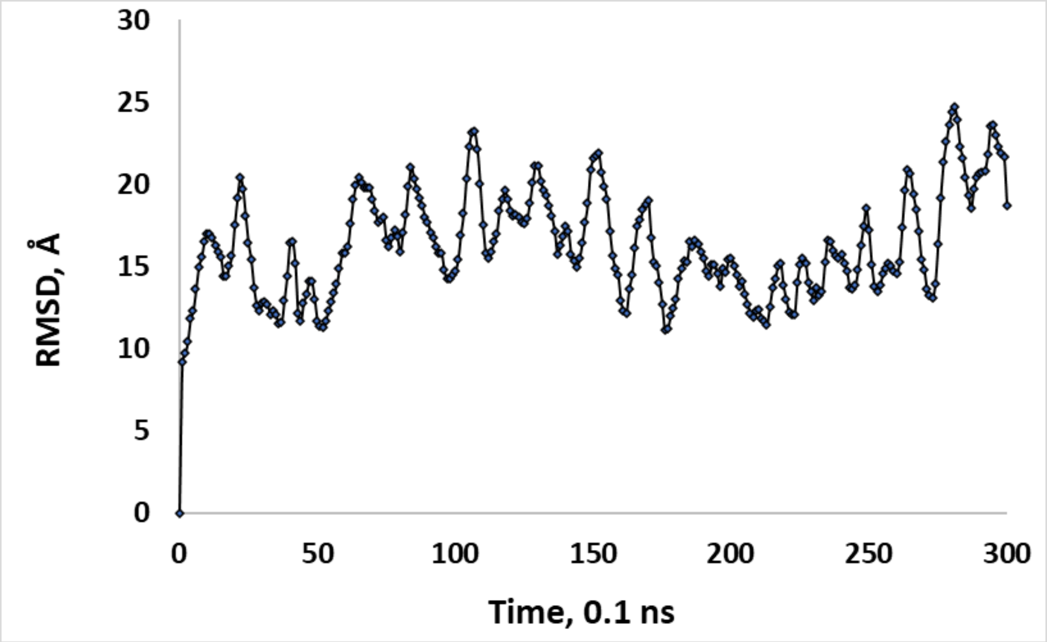


**Figure S20**. RMSD plot of the pre-designed spiral with lateral interactions inside the 40-nm tube (Fig S8)
